## Supplementary material for "A system for functional studies of the major virulence factor of malaria parasites": Supplmental figures

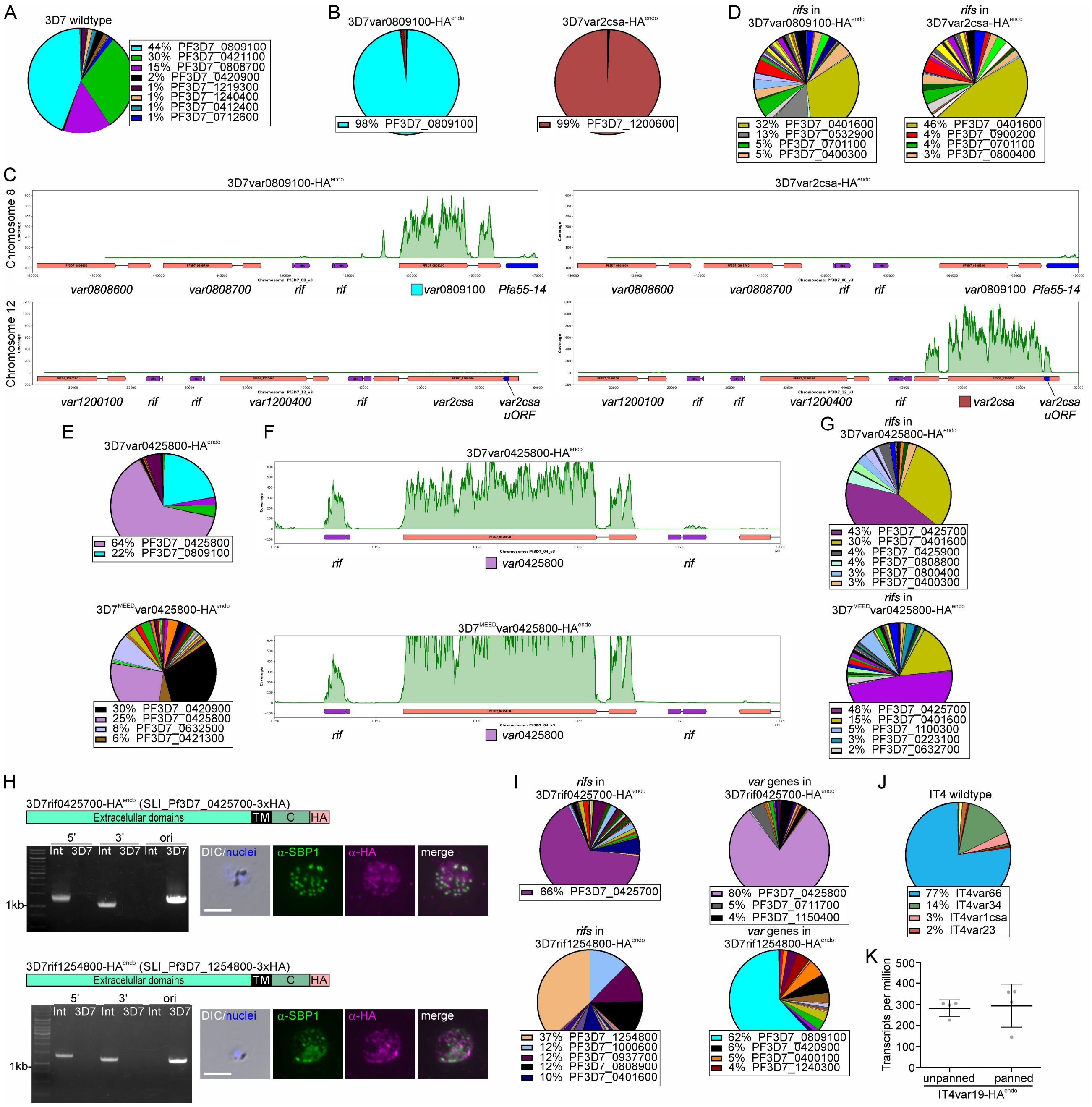


**Figure S1. Expression of desired and co-activated genes.** (**A**) Pie chart with proportions of total *var* gene transcripts determined by qPCR of the 3D7 parent. (**B**) Pie charts with proportions of total *var* gene transcripts of the indicated cell lines determined by RNAseq (normalized to TPM) showing predominant expression of the desired SLI-activated *var* gene. (**C**) Coverage plots showing reads mapped to chromosome 8 base pairs 430000 to 470000 and chromosome 12 base pairs 15000 to 60000 of the indicated cell lines determined by RNAseq (shows reciprocal activity of *var* genes as expected based on the SLI selected target as well as inactivity of neighbouring *var* genes). Gene annotation from PlasmoDB genome browser (PlasmoDB.org) with reads mapped using Artemis. Red: *var* genes; purple: *rif* genes. Colored boxes indicated SLI activated *var* gene. (**D**) Pie charts with proportions of total *rif* gene transcripts of the indicated SLI-activated *var* gene cell lines determined by RNAseq (normalized to TPM). (**E**) Pie charts with proportions of total *var* gene transcripts of the indicated cell lines determined by qPCR. (**F**) Coverage plots showing mapped reads on chromosome 4 base pairs 1150000 to 1175000 of the indicated cell lines determined by RNAseq as in (C). Coloured boxes indicate SLI activated *var* gene. Red: *var* genes; purple: *rif* genes. (**G**) Pie charts with proportions of total *rif* gene transcripts of the indicated cell lines determined by RNAseq (normalized to TPM). (**H**) Confirmation of the activation of the indicated *rif* gene. Scheme shows domain organisation (TM, transmembrane domain; C, C-terminal domain; HA, 3xHA tag). Agarose gel shows PCR products confirming correct integration of the SLI plasmid as described in Fig. 1A; 3D7 parent; Int: integrant cell line. Fluorescence microscopy: images of IFAs with the indicated antibodies. Nuclei: Hoechst 33342; DIC: differential interference contrast; size bars 5 µm. (**I**) Pie charts show proportions of total *var* or *rif* gene transcripts of the indicated cell lines determined by qPCR. (**J**) Pie charts with proportions of total *var* gene transcripts of IT4 wildtype parasites determined by RNAseq (normalized to TPM). (**K**) Plot showing transcription levels of *var19* in the IT4var19-HA$^{endo}$ parasites before and after panning determined by RNAseq (normalized to TPM) (n = 4).

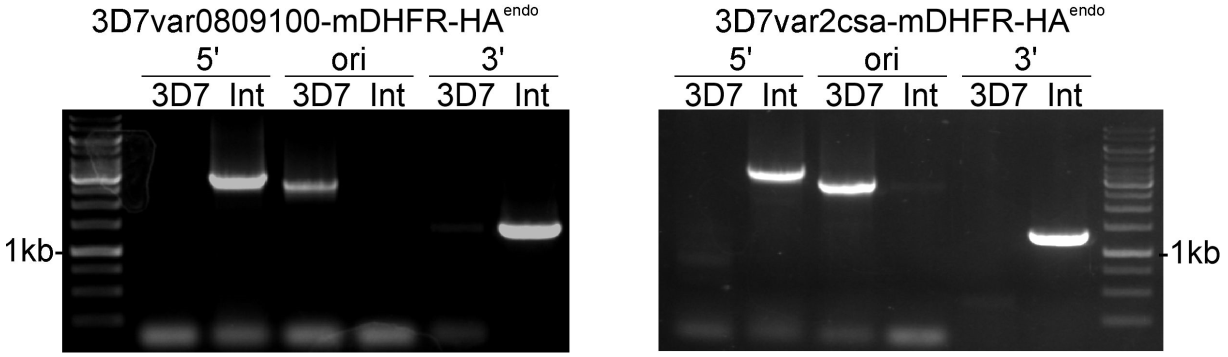


**Figure S2. Confirmation of correct integration for PfEMP1 mDHFR fusion parasites.** Agarose gels with PCR products confirming correct integration of the SLI plasmids to generate the indicated cell lines. Product over 5′ integration junction (5'): P1+P2; 3' integration junction (3'): P3+P4; original locus (ori): P1+P4; see Fig. 1A for primer positions, Table S6 for primer sequences; 3D7: parent; Int: integrant cell line.

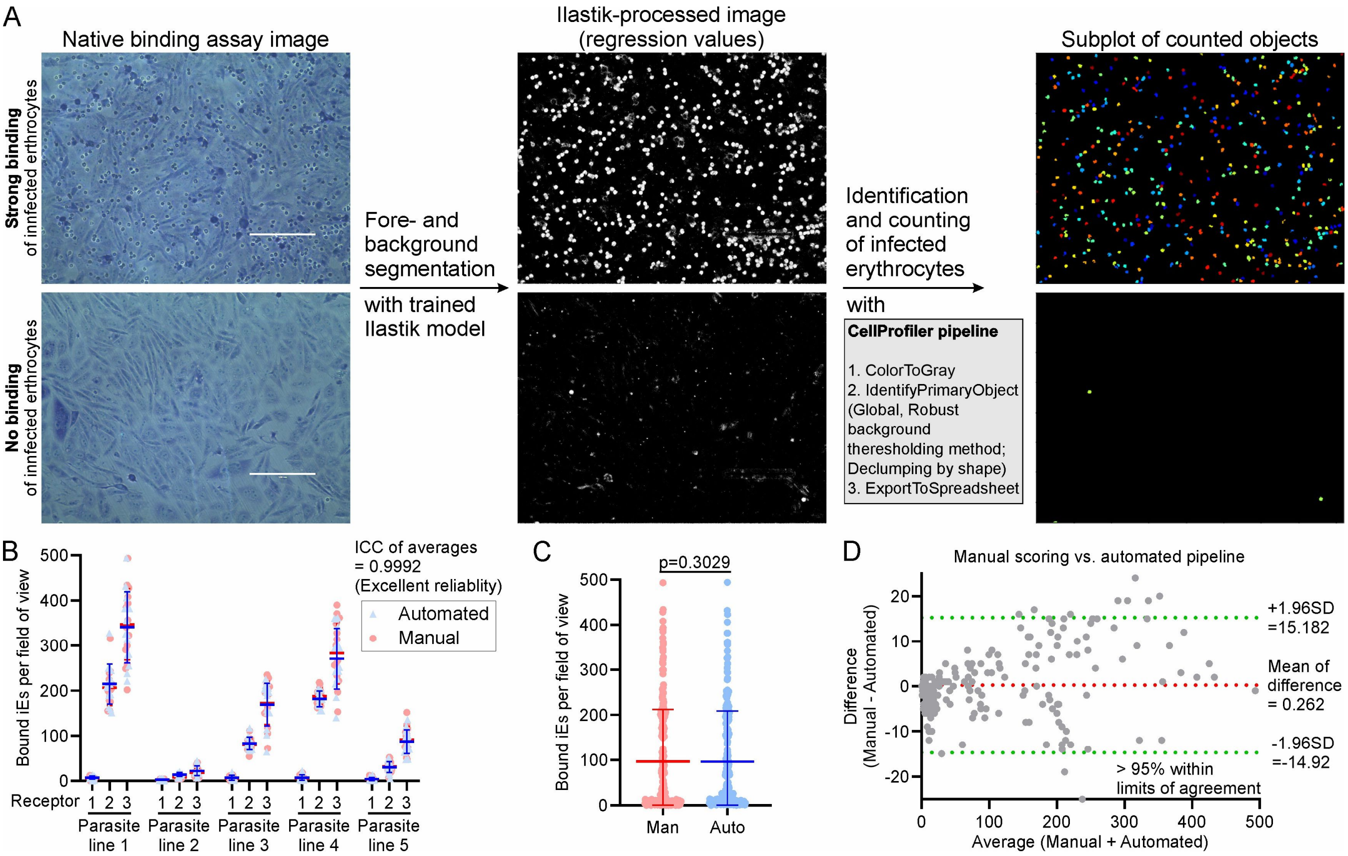


**Figure S3. Automated scoring pipeline with validation.** (**A**) Illustration of the pipeline for automated scoring of the number of bound infected erythrocytes in images captured from binding assays. Left: representative image of a binding assay showing binding (top) or no binding (bottom) of infected erythrocytes. Ilastik [119] model (trained on 20 images) separates fore- and background of native captured images (left, input images; middle, output images). Foreground: infected erythrocytes; background: CHO/HBEC-5i cells and plastic. Gray box: CellProfiler [120] pipeline to score pre-segmented images (middle, input images; right, output images). (**B**) Results of binding assays for five parasite lines tested against three different receptor expressing CHO-cells evaluated by manual scoring and the automated pipeline (15 fields of views were analysed per cell line and receptor, total: 225; bars: mean and SD). ICC: intraclass correlation coefficient. (**C**) Comparison of all images from (B) evaluated by manual scoring against scoring by the automated pipeline. Red dots: bound infected erythrocytes (iE) in individual images from manual scoring (blue dots) or scoring by the automated pipeline. Man: manual scoring; auto: automated pipeline (n = 225; bars: mean and SD; paired t-test; p-value is indicated). (**D**) Bland-Altman plot comparing evaluation of images of binding assays from (B) by manual scoring and scoring by the automated pipeline. Green lines: limits of agreement; SD: standard deviation.

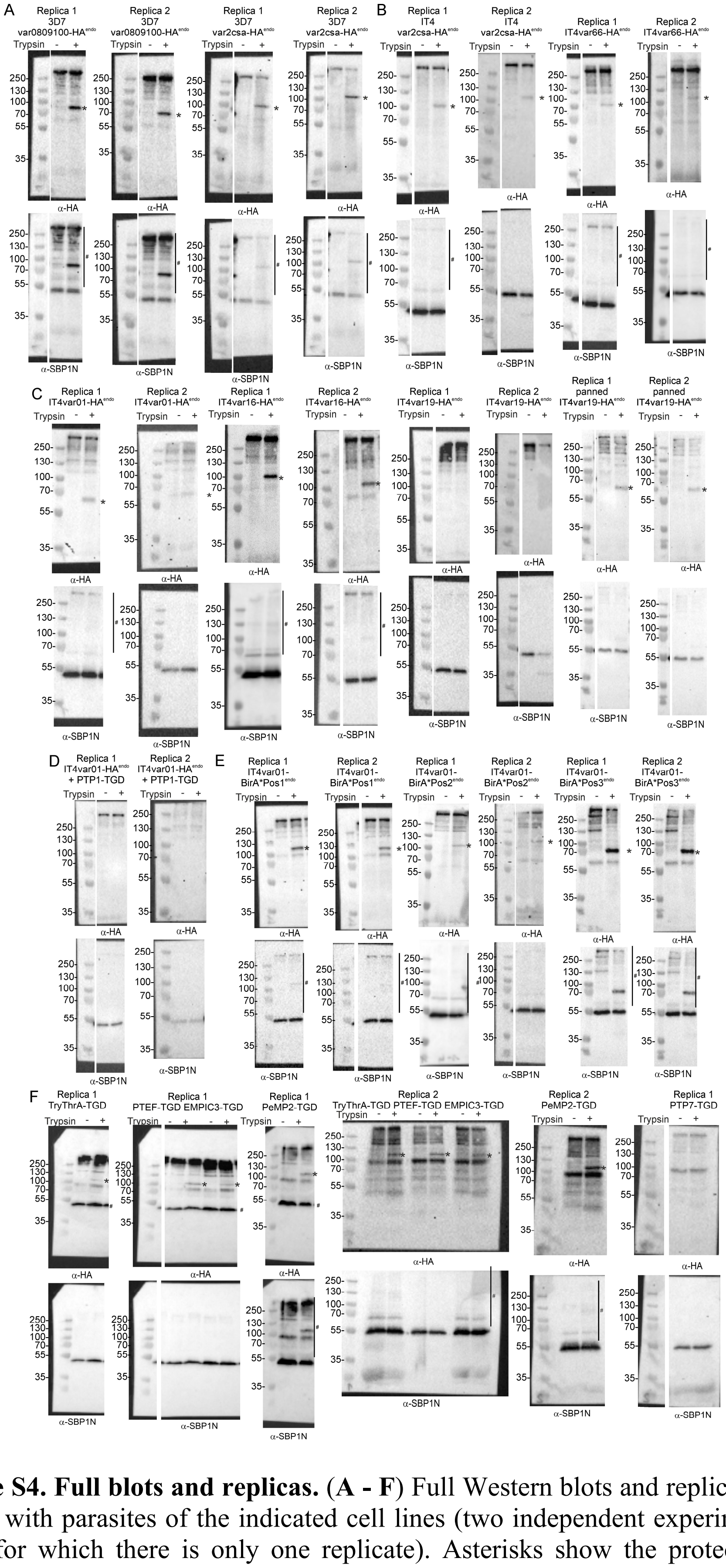

**Figure S4. Full blots and replicas.** (**A - F**) Full Western blots and replicates of trypsin cleavage assays with parasites of the indicated cell lines (two independent experiments except for PTP7-TGD for which there is only one replicate). Asterisks show the protected PfEMP1 fragment indicative of surface exposure. Hash sign indicates signal from previous probing of the blot. α-SBP1-N: control for integrity of host cell (breach of RBC membrane would result in a smaller SBP1 fragment). Marker in kDa.

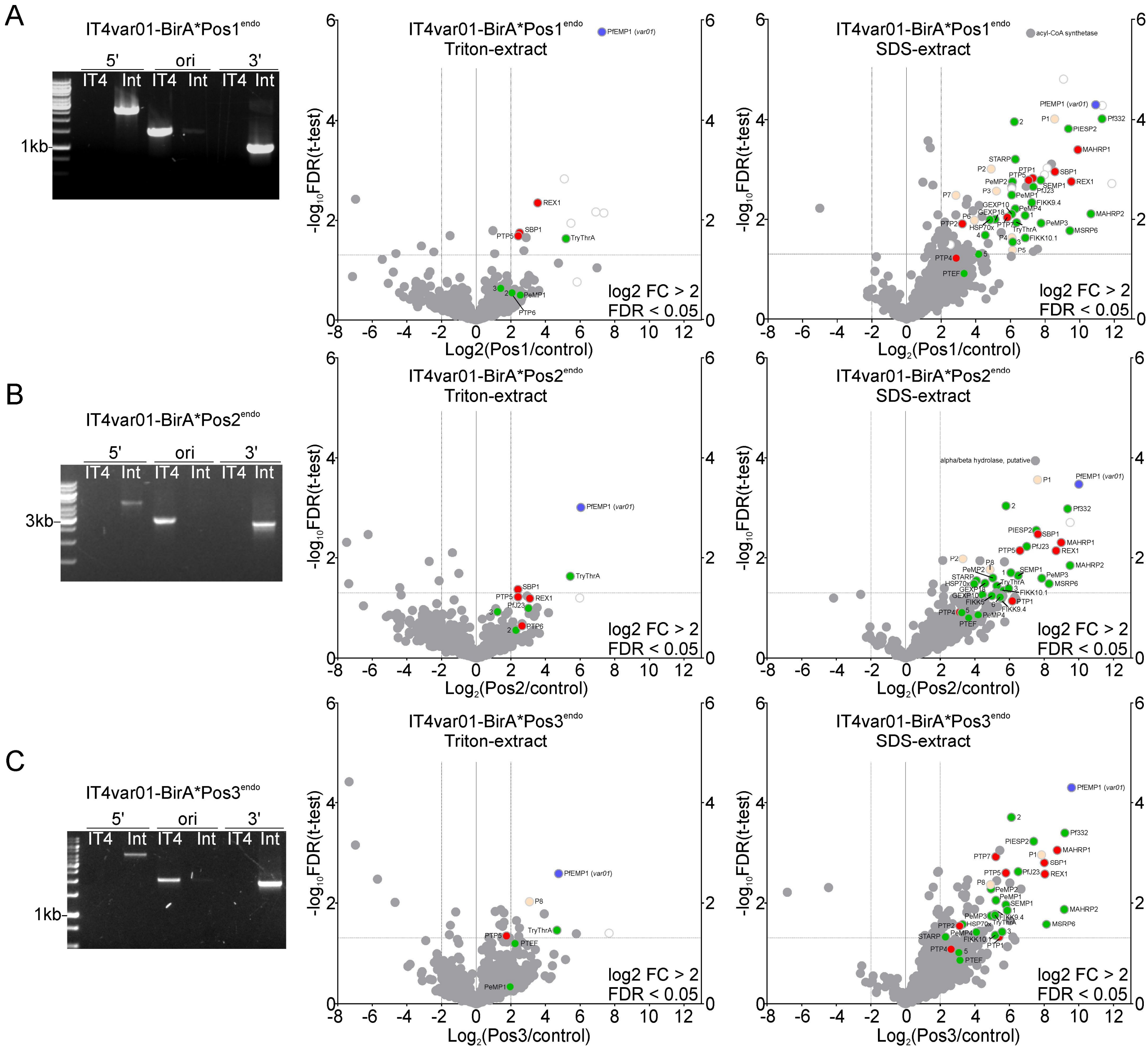


**Figure S5. Correct integration of BioID cell lines and full volcano plots.** (**A**, **B**, **C**) Agarose gels show PCR products confirming correct integration of the SLI plasmid for the indicated cell lines. Product over 5′ integration junction (5'): P1+P2; 3' integration junction (3'): P3+P4; original locus (ori): P1+P4; see Fig. 1A for primer positions, Table S6 for sequence; IT4: parent; Int: integrant cell line. Graphs: full volcano plots showing enrichment of biotinylated proteins extracted with Triton (middle row) or SDS (right row, full plots of plots in Fig. 6) from the lysates of the indicated cell lines compared to IT4 wildtype parasites. Confidence above – $\log_{10}$ FDR of 0.05 and $\log_2$ enrichment of 2 are indicated by red lines (Full data in Table S4).

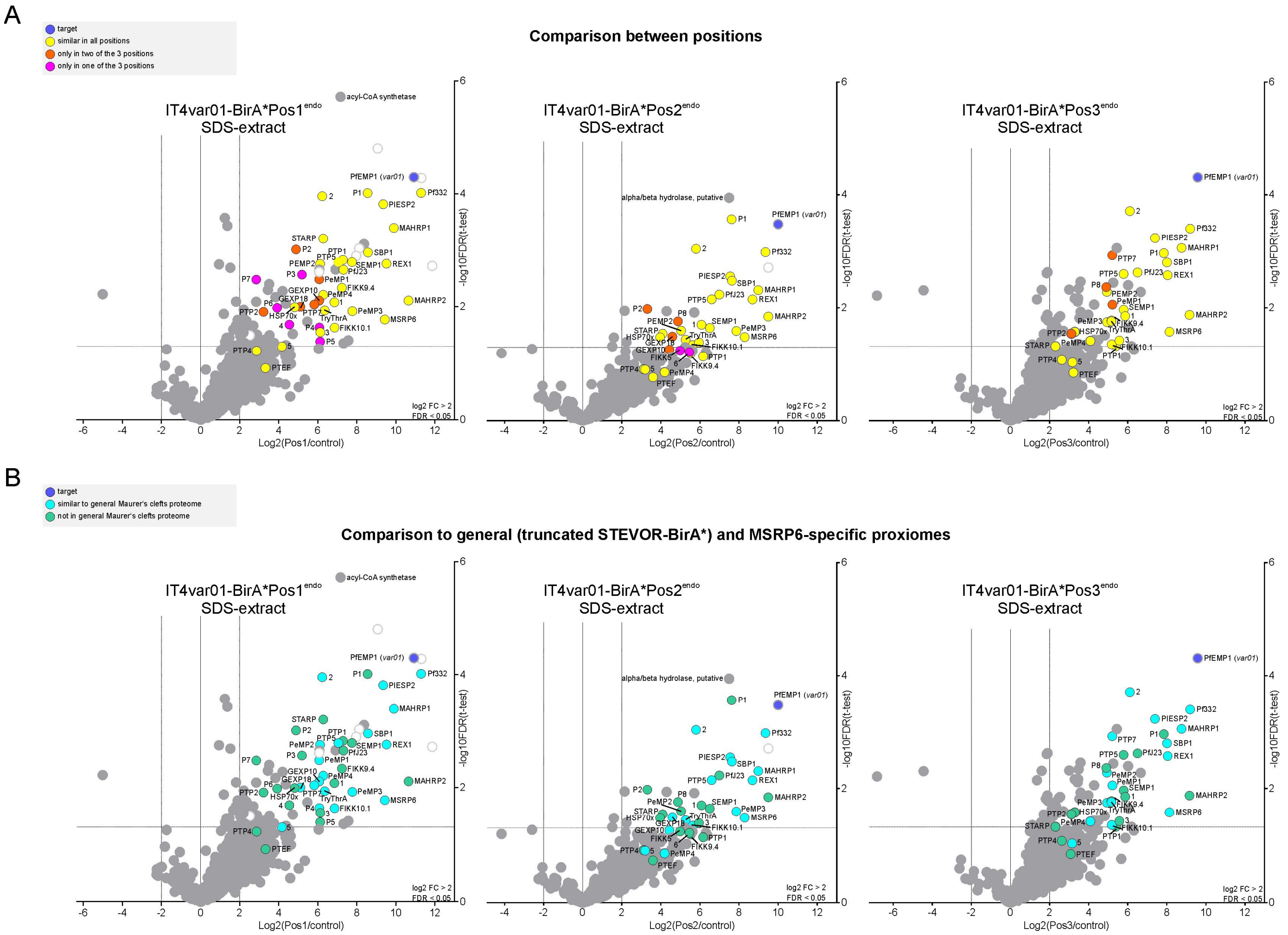


**Figure S6. Volcano plots with colour coding for comparison.** (**A**, **B**) Volcano plots of SDS extracts shown in Fig. S5 with colour coding according to similarity between positions (A) or with a general Maurer´s cleft BioID [67] (B). Hits were considered similar in all positions (yellow) when they were in a similar relative position to other hits (A). Hits were in (B) marked as present in general Maurer’s clefts proteome (light blue) if significantly enriched in BioID experiments of a general Maurer’s clefts marker over a protein soluble in the host cell or in the Maurer’s clefts attachment domain of MSRP6 over the control protein soluble in the host cell [67].

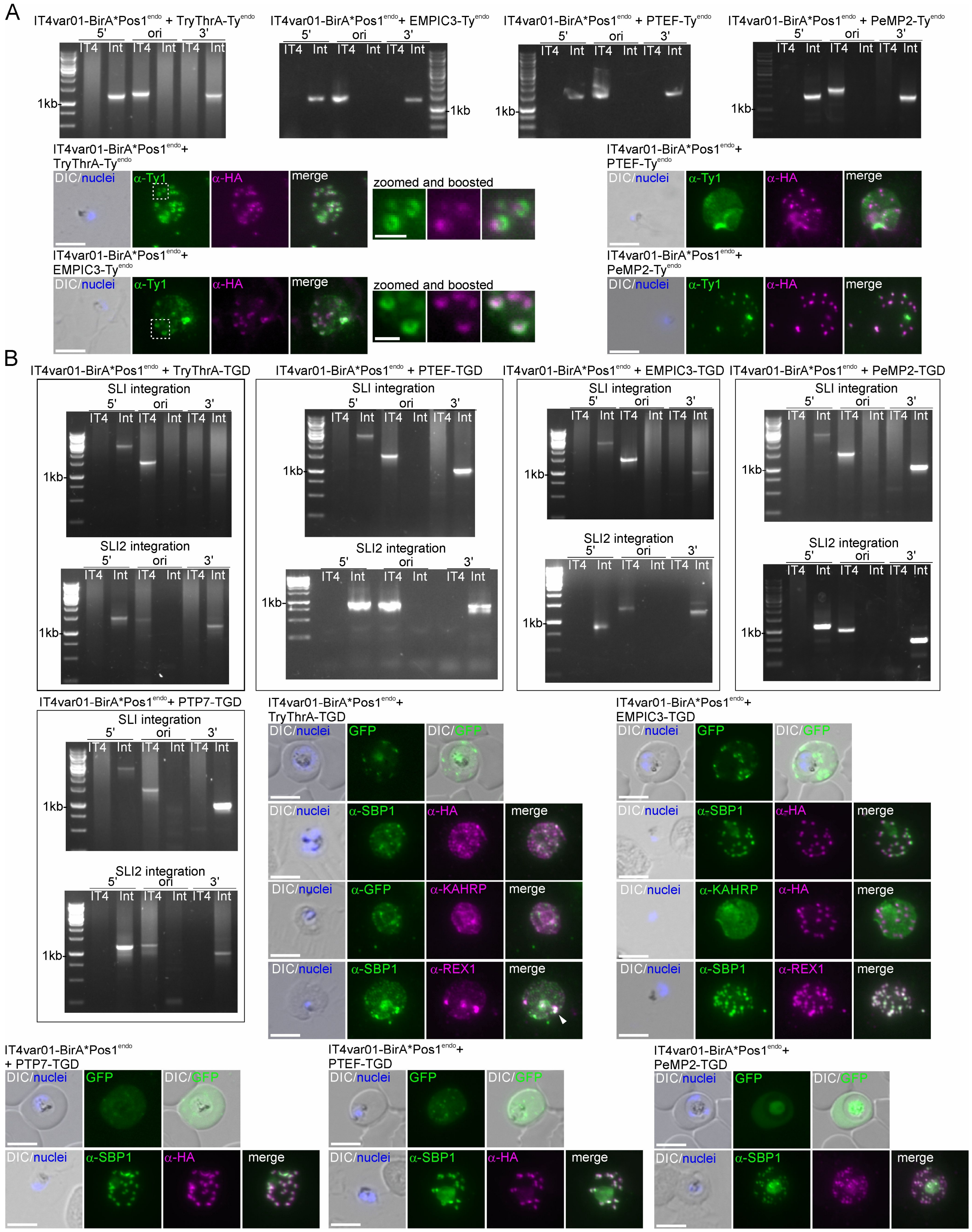


**Figure S7. Cell lines and analysis of cytoadherence protein candidates.** (**A**) Agarose gels show PCR products confirming correct integration of the SLI2 plasmid for the indicated cell line. PCR over 5´integration junction (5'): P5 + P8 in Fig. 5A; PCR over 3' integration junction (3'): P7 +

P6 in Fig. 5A; original locus (ori): P5 + P6 in Fig. 5A; IT4 parent; Int: integrant cell line; primers in Table S6. Fluorescence microscopy images show IFAs of the indicated cell lines with indicated antibodies. Zoomed and boosted: enlargements with boosted intensity of the boxed areas to illustrate the circular signal of TryThrA-Ty and EMPIC3-Ty. Nuclei: Hoechst 33342; DIC: differential interference contrast; size bars 5 µm and 1 µm in the zoomed images. (**B**) Agarose gels show PCR products confirming correct integration of the SLI2 plasmid and perpetuation of SLI plasmid integration for the indicated cell lines. PCR over 5´integration junction (5'): P1 + P2 in Fig. 1A or P5 + P8 in Fig. 5A; PCR over 3' integration junction (3'): P3 + P4 in Fig. 1A or P7 + P6 in Fig. 5A; original locus (ori): P1 + P4 in Fig. 1A or P5 + P6 in Fig. 5A; IT4 parent; Int: integrant cell line; primers in Table S6. Fluorescence microscopy images show live parasites (top rows) or IFAs (bottom rows) with indicated antibodies of the TGD cell lines. Nuclei: Hoechst 33342; DIC: differential interference contrast; size bars 5 µm.

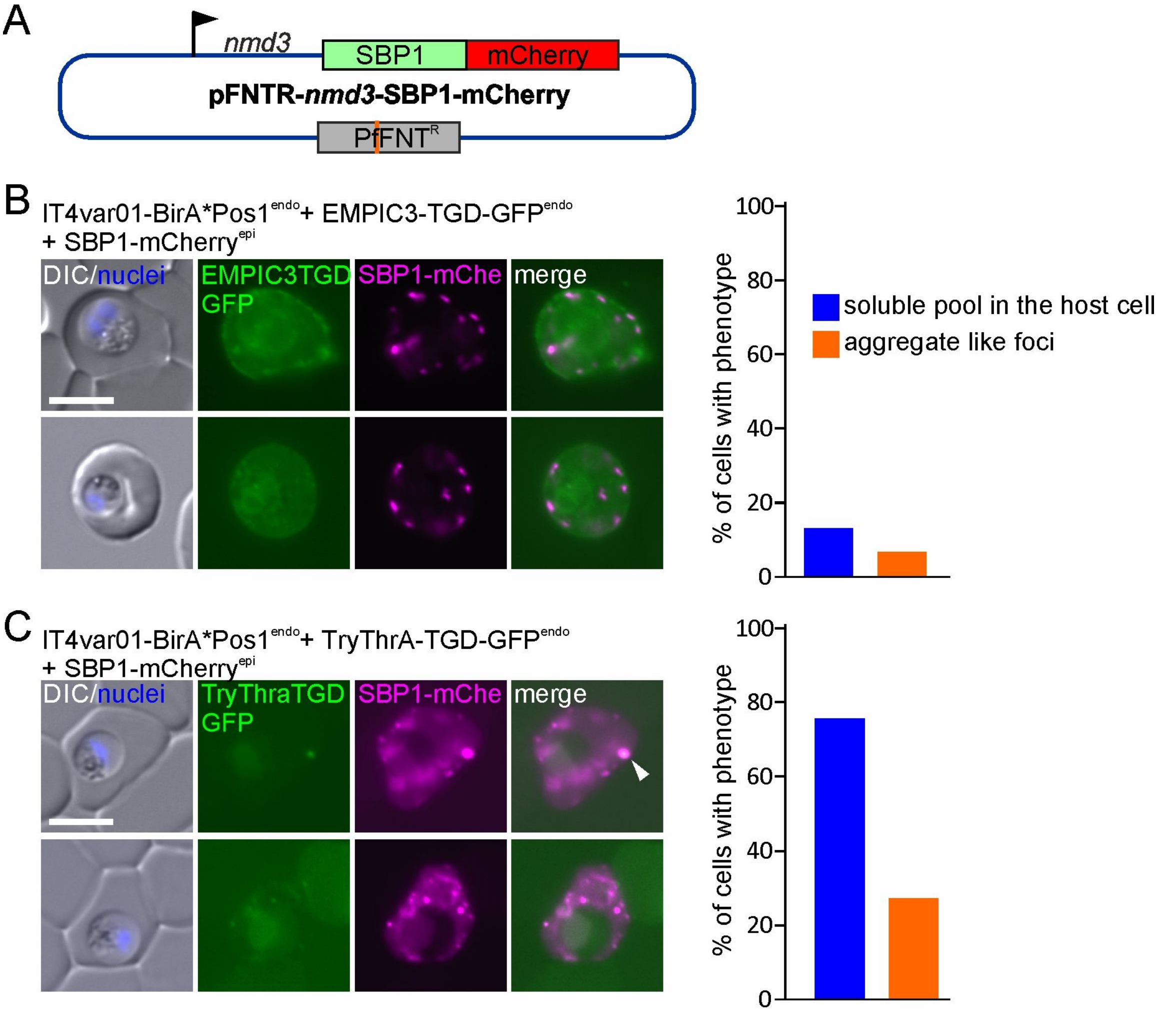


**Figure S8. Live cell imaging of episomally expressed SBP1-mCherry in the EMPIC3- and TryThrA-TGD parasites.** (**A**) Scheme of plasmid used to episomally express SBP1-mCherry in the parasites containing already two SLI modifications (in this case the IT4var01-BirA*Pos1$^{endo}$ cell line with SLI2-mediated TGDs as indicated in B and C). The orange bar indicates the position of the G 107 S change in PfFNT conferring resistance to BH267.meta. **(B)** and **(C)**, Live cell fluorescence microscopy images of the indicated cell lines containing the plasmid shown in (A) (left) and quantification of phenotype (right; data from 31 cells from 7 independent imaging sessions for the EMPIC3-TGD and 37 cells from 7 independent sessions for the TryThrA-TGD). The arrow shows what was scored as aggregate (disproportionally strong focus compared to other foci). The truncated EMPIC3 and TryThrA are fused with GFP. SBP1-mChe, mCherry fused SBP1; Nuclei: Hoechst 33342; DIC: differential interference contrast; size bars 5 µm.
