## Supplementary material for "A system for functional studies of the major virulence factor of malaria parasites": Data S1 (plasmids)

SLI integration constructs (3xHA)

XXXXX = hDHFR (WR resistance)

XXXXX = 3xHA

XXXXX = Skip peptide

XXXXX = Neomycin

>pSLI-PfIT_060021400-3xHA-T2A-NeoR

ggatatggcagcttaatgttcgtttttcttatttatatatttataccaattgattgtatttataactgtaaaaatgtgtatgttgtgtgcatatttttttttgtgcatgcacatgcatgtaaatagctaaaattatgaacattttattttttgttcagaaaaaaaaaactttacacacataaaatggctagtatgaatagccatattttatataaattaaatcctatgaatttatgaccatattaaaaatttagatatttatggaacataatatgtttgaaacaataagacaaaattattattattattattatttttactgttataattatgtgtctccttcaatgattcataaatagttggacttgatttttaaaatgtttataatatgattagcatagttaaataaaaaaagttgaaaaattaaaaaaaaacatataaacacaaatgatggtttttccttcaatttcgatatcaatttatagaaacaaaatatatacttgtataattttatttttttatataaatcattacatatataattatacaatattttttctaagagataattatatattaatatatataaaaaaaggtgttttttttttttttttttatttttatttttattttatggtaatattttattttccttattttataaattatattagtttatatgtgattaattttatatattatcaatttatatatttttaaatgcttacttaattatctttttttttttttttttttttttttcccctctttttatattaatttatttttgaaaaaattgatatatatatatatatataatatatatatatacatgtagtagtattaaacaatgtataatatatataaataatatatttatatatttcatttcaattttaattttttttggttttttttttttttctttttgtcatatttaaaaaaaattatattcatataagttatgcattttttataaacattattcaatatatgtataatataatatatatatatatattaatgtattattccaatgtgcatgataaaagaaaaaaataatatttataaaaaaaaagaaaaataaaacaaaaaaagaaaaaaaaaaaaaaaaaaaaaaaaatacaaaaataaataatataatttataattatatattcttgtcacaataaaaatatatatatatatatatatatttataatatgtatattttaaactagaaaaggaataactaatattttatttattatcattcaagatttatattttataataataaatacctaatagaaatatatcaggatccatgcatggttcgctaaactgcatcgtcgctgtgtcccagaacatgggcatcggcaagaacggggactacccctggccaccgctcaggaacgaatttagatatttccagagaatgaccacaacctcttcagtagaaggtaaacagaatctggtgattatgggtaagaagacctggttctccattcctgagaagaatcgacctttaaagggtagaattaatttagttctcagcagagaactcaaggaacctccacaaggagctcattttctttccagaagtctagatgatgccttaaaacttactgaacaaccagaattagcaaataaagtagacatggtctggatagttggtggcagttctgtttataaggaagccatgaatcacccaggccatcttaaactatttgtgacaaggatcatgcaagactttgaaagtgacacgttttttccagaaattgatttggagaaatataaacttctgccagaatacccaggtgttctctctgatgtccaggaggagaaaggcattaagtacaaatttgaagtatatgagaagaatgattaagcttatttaataatagattaaaaatattataaaaataaaaacataaacacagaaattacaaaaaaaatacatatgaattttttttttgtaatcttccttataaatatagaataatgaatcatataaaacatatcattattcatttatttacatttaaaattattgtttcagtatctttaatttattatgtatatataaaaataacttacaattttattaataaacaatatatgtttattaattcatgttttgtaatttatgggatagcgattttttttactgtctgtatttttcttttttaattatgttttaattgtattttatttttattattgttctttttatagtattattttaaaacaaaatgtattttctaagaacttataataataataaatataaattttaataaaaattatatttatcttttacaatatgaacataaagtacaacattaatatatagcttttaatatttttattcctaatcatgtaaatcttaaatttttctttttaaacatatgttaaatatttatttctcattatatataagaacatatttattaaatctagaattctatagtgagtcgtattacaattcactggccgtcgttttacaacgtcgtgactgggaaaaccctggcgttacccaacttaatcgccttgcagcacatccccctttcgccagctggcgtaatagcgaagaggcccgcaccgatcgcccttcccaacagttgcgcagcctgaatggcgaatggcgcctgatgcggtattttctccttacgcatctgtgcggtatttcacaccgcatatggtgcactctcagtacaatctgctctgatgccgcatagttaagccagccccgacacccgccaacacccgctgacgcgccctgacgggcttgtctgctcccggcatccgcttacagacaagctgtgaccgtctccgggagctgcatgtgtcagaggttttcaccgtcatcaccgaaacgcgcgagacgaaagggcctcgtgatacgcctatttttataggttaatgtcatgataataatggtttcttagacgtcaggtggcacttttcggggaaatgtgcgcggaacccctatttgtttatttttctaaatacattcaaatatgtatccgctcatgagacaataaccctgataaatgcttcaataatattgaaaaaggaagagtatgagtattcaacatttccgtgtcgcccttattcccttttttgcggcattttgccttcctgtttttgctcacccagaaacgctggtgaaagtaaaagatgctgaagatcagttgggtgcacgagtgggttacatcgaactggatctcaacagcggtaagatccttgagagttttcgccccgaagaacgttttccaatgatgagcacttttaaagttctgctatgtggcgcggtattatcccgtattgacgccgggcaagagcaactcggtcgccgcatacactattctcagaatgacttggttgagtactcaccagtcacagaaaagcatcttacggatggcatgacagtaagagaattatgcagtgctgccataaccatgagtgataacactgcggccaacttacttctgacaacgatcggaggaccgaaggagctaaccgcttttttgcacaacatgggggatcatgtaactcgccttgatcgttgggaaccggagctgaatgaagccataccaaacgacgagcgtgacaccacgatgcctgtagcaatgccaacaacgttgcgcaaactattaactggcgaactacttactctagcttcccggcaacaattaatagactggatggaggcggataaagttgcaggaccacttctgcgctcggcccttccggctggctggtttattgctgataaatctggagccggtgagcgtgggtctcgcggtatcattgcagcactggggccagatggtaagccctcccgtatcgtagttatctacacgacggggagtcaggcaactatggatgaacgaaatagacagatcgctgagataggtgcctcactgattaagcattggtaactgtcagaccaagtttactcatatatactttagattgatttaaaacttcatttttaatttaaaaggatctaggtgaagatcctttttgataatctcatgaccaaaatcccttaacgtgagttttcgttccactgagcgtcagaccccgtagaaaagatcaaaggatcttcttgagatcctttttttctgcgcgtaatctgctgcttgcaaacaaaaaaaccaccgctaccagcggtggtttgtttgccggatcaagagctaccaactctttttccgaaggtaactggcttcagcagagcgcagataccaaatactgtccttctagtgtagccgtagttaggccaccacttcaagaactctgtagcaccgcctacatacctcgctctgctaatcctgttaccagtggctgctgccagtggcgataagtcgtgtcttaccgggttggactcaagacgatagttaccggataaggcgcagcggtcgggctgaacggggggttcgtgcacacagcccagcttggagcgaacgacctacaccgaactgagatacctacagcgtgagctatgagaaagcgccacgcttcccgaagggagaaaggcggacaggtatccggtaagcggcagggtcggaacaggagagcgcacgagggagcttccagggggaaacgcctggtatctttatagtcctgtcgggtttcgccacctctgacttgagcgtcgatttttgtgatgctcgtcaggggggcggagcctatcgaaaaacgccagcaacgcggcctttttacggttcctggccttttgctggccttttgctcacatgttctttcctgcgttatcccctgattctgtggataaccgtattaccgcctttgagtgagctgataccgctcgccgcagccgaacgaccgagcgcagcgagtcagtgagcgaggaagcggaagagcgcccaatacgcaaaccgcctctccccgcgcgttggccgattcattaatgcagctggcacgacaggtttcccgactggaaagcgggcagtgagcgcaacgcaattaatgtgagttagctcactcattaggcaccccaggctttacactttatgcttccggctcgtatgttgtgtggaattgtgagcggataacaatttcacacaggaaacagctatgaccatgattacgccaagctatttaggtgacactatagaatactc**gcggccgcTAAINSERTGGTACC**ACCATGTACCCATACGATGTTCCAGATTACGCTACGATGTACCCGTACGACGTGCCGGACTACGCGACTATGTATCCATATGATGTTCCAGATTATGCTGTCGACGGAGAAGGAAGAGGAAGTTTATTAACATGTGGAGATGTAGAAGAAAATCCAGGACCAATGATTGAACAAGATGGATTGCACGCAGGTTCTCCGGCCGCTTGGGTGGAGAGGCTATTCGGCTATGACTGGGCACAACAGACAATCGGCTGCTCTGATGCCGCCGTGTTCCGGCTGTCAGCGCAGGGGCGCCCGGTTCTTTTTGTCAAGACCGACCTGTCCGGTGCCCTGAATGAACTGCAGGACGAGGCAGCGCGGCTATCGTGGCTGGCCACGACGGGCGTTCCTTGCGCAGCTGTGCTCGACGTTGTCACTGAAGCGGGAAGGGACTGGCTGCTATTGGGCGAAGTGCCGGGGCAGGATCTCCTGTCATCTCACCTTGCTCCTGCCGAGAAAGTATCCATCATGGCTGATGCAATGCGGCGGCTGCATACGCTTGATCCGGCTACCTGCCCATTCGACCACCAAGCGAAACATCGCATCGAGCGAGCACGTACTCGGATGGAAGCCGGTCTTGTCGATCAGGATGATCTGGACGAAGAGCATCAGGGGCTCGCGCCAGCCGAACTGTTCGCCAGGCTCAAGGCGCGCATGCCCGACGGCGAGGATCTCGTCGTGACCCATGGCGATGCCTGCTTGCCGAATATCATGGTGGAAAATGGCCGCTTTTCTGGATTCATCGACTGTGGCCGGCTGGGTGTGGCGGACCGCTATCAGGACATAGCGTTGGCTACCCGTGATATTGCTGAAGAGCTTGGCGGCGAATGGGCTGACCGCTTCCTCGTGCTTTACGGTATCGCCGCTCCCGATTCGCAGCGCATCGCCTTCTATCGCCTTCTTGACGAGTTCTTCTAACTCGAG

Inserts:

>3D7var0809100

ACCAGGTAGTCCTAAATATAAAACGTTGATAGAAGTGGTACTTGAACCATCAAAAAGAGATATACCAAGTGGTGATATACCACATACAAATAAATTTACAGATAATGAATGGAATCAATTGAAAAAAGATTTTATATCTAATATGTTACAAAATACCCAAAATACGGAACCAAATGTTTTACATGATAATGTGGATAATAATACCCATCCTACCATGTCACGTCATAATGTGGACCAAAAACCTTTTATTATGTCCATACATGATAGAAATTTATATATTGGAGAAGAATATAGTTATGATATGAGTACTAACAGTGGTGAAAATAATTTATACAGTGGTATTGATCCAACAAGTGCTAACCATGATTCTTATAGTGGTATAGATTTAATCAATGACGCACTAAATGGTGATTATGACATTTACGATGAAATATTGAAACGAAAAGAAAATGAATTATTTGGGACAAATCATACGAAAAAAAATACATCAACCAATAGTGTTGCAAAAAATACAAATAGTGATCCTATACTCAATCAAATAAATTTGTTCCATAAATGGTTAGATAGACATAGAAATATGTGCGAACAGTGGGATAAAAATAAAAAGGAGGAATTTTTAGATAAATTAAAAAAAGAATGGAACAAAGAAAATAACAATAATAGTGGTGACATTAACAATAGGTATGAGAATGTGTTGAATACTGATGTTTCTATTCAAATAGATATGCATAATCCTAAACCAAAGAACGAATTTACGAATATGGATACAAACCCTGATAATTTCATTAAGGATACTATATTGAATGATCTGGAAAAACATCGTGAACCCTACTTCTATGATATTTATGATGATGATATCACTTATTTTGATACAGATGATGTTAAACCACCTATGGATGATATACACATTAAAGAACAAACTGAAATGAATGCTCTTCATAATAATAAGATGAATGAATTGTTAGAAAAGGAATATCCTATATCAGATATATGGAATATA

>3D7var2csa

CGATAAATATATTTGGGACTTATCTTCCTCTGATATAACTTCCTCCGAAAGTGAGTATGAAGAAGTGGATATCAATGATATATATGTACCAAGTTTTCCCAAATATAAAACGTTCATTGAATTAGTACTAGAACCTTCCAAAAGGGATACATTTAATACATCAAGTGGTGACACATTCACCAATAAACTTACGGATGATGAATGGAACCAATTGAAACAGGATTTTATTGAACAATATTTACAAAACATACAAAAGGATTTTATTTTACATGATAGTATGGATGAAAAACCTTTTATTACTCAAATCCAGGATAGATTTCTTGATAGTAGTCATGAAGAAGTTACTTATAATATTGATTGGAATGTTCCTGAAAATATTAATAGGATTACTAATAACATGGACGATCCAAAATACTGCTCAAATAATATGTATACTGGTACCGATTTAATTAATGATTCATTAAATGGTAACCAATATATTGATATATATGATGAGATGCTGAAACGAAAAGAAAACGAATTATTTGGAACATATCATACAAAATATACAACCTTTAACAGTGTTTCTAAACAAACACCTAGTGACCCGATAATTAACCAACTAGATTTATATCATAAATGGATAGACAAACATAGAGATATTTGCGAACAGTGGAAAACCAAAGAGGATATGTTATATAAATTGAATGAAGTGTGGAATATGGAACGTAAGGAATATCTATTGGATATACAACCATCAACTCTGGATGATATTCATAAAATTAATGATGAAACATATAATATTATTAGTACAAATAATATATATGATCATCCCTCACAGGAAACCCCCCTCCAACTACTTGGATCAACAAATATTATACCCAGTTATATTACCACGGAACAAAATAATGGATTGCGCACAAATATATCTATGGATACATATATTGATGAAACAAATAATAATAATGTGGTAGCCACTAGTATAATAGGTGACGATCAGATGGAAAATTCGTACAATTGT

>3D70809100-mDHFR

ACCAGGTAGTCCTAAATATAAAACGTTGATAGAAGTGGTACTTGAACCATCAAAAAGAGATATACCAAGTGGTGATATACCACATACAAATAAATTTACAGATAATGAATGGAATCAATTGAAAAAAGATTTTATATCTAATATGTTACAAAATACCCAAAATACGGAACCAAATGTTTTACATGATAATGTGGATAATAATACCCATCCTACCATGTCACGTCATAATGTGGACCAAAAACCTTTTATTATGTCCATACATGATAGAAATTTATATATTGGAGAAGAATATAGTTATGATATGAGTACTAACAGTGGTGAAAATAATTTATACAGTGGTATTGATCCAACAAGTGCTAACCATGATTCTTATAGTGGTATAGATTTAATCAATGACGCACTAAATGGTGATTATGACATTTACGATGAAATATTGAAACGAAAAGAAAATGAATTATTTGGGACAAATCATACGAAAAAAAATACATCAACCAATAGTGTTGCAAAAAATACAAATAGTGATCCTATACTCAATCAAATAAATTTGTTCCATAAATGGTTAGATAGACATAGAAATATGTGCGAACAGTGGGATAAAAATAAAAAGGAGGAATTTTTAGATAAATTAAAAAAAGAATGGAACAAAGAAAATAACAATAATAGTGGTGACATTAACAATAGGTATGAGAATGTGTTGAATACTGATGTTTCTATTCAAATAGATATGCATAATCCTAAACCAAAGAACGAATTTACGAATATGGATACAAACCCTGATAATTTCATTAAGGATACTATATTGAATGATCTGGAAAAACATCGTGAACCCTACTTCTATGATATTTATGATGATGATATCACTTATTTTGATACAGATGATGTTAAACCACCTATGGATGATATACACATTAAAGAACAAACTGAAATGAATGCTCTTCATAATAATAAGATGAATGAATTGTTAGAAAAGGAATATCCTATATCAGATATATGGAATATACCCGGGATGGTAAGGCCTTTGAATTGTATAGTTGCAGTTAGTCAAAATATGGGAATTGGGAAAAATGGTGATCTTCCATGGCCCCCCTTACGTAATGAGTTCAAGTACTTTCAAAGGATGACTACCACAAGTAGTGTGGAGGGAAAGCAAAACTTAGTAATAATGGGGCGTAAAACTTGGTTTAGTATACCCGAAAAAAATAGGCCATTGAAAGATCGTATAAACATTGTATTGTCACGTGAGTTGAAAGAGCCACCTAGAGGAGCACACTTCTTGGCTAAATCTTTGGACGACGCTTTGAGGTTAATAGAGCAGCCTGAGCTTGCTTCAAAGGTTGATATGGTATGGATTGTAGGAGGTTCAAGTGTATACCAAGAGGCAATGAACCAACCTGGACACTTAAGGTTGTTCGTAACTCGTATTATGCAGGAGTTCGAGTCAGATACATTCTTCCCTGAGATAGACCTTGGAAAGTACAAGTTGTTACCTGAGTATCCTGGAGTATTAAGTGAAGTTCAAGAAGAAAAGGGAATAAAATATAAGTTCGAGGTTTACGAAAAAAAGGAC

>3D7var2csa-mDHFR

CGATAAATATATTTGGGACTTATCTTCCTCTGATATAACTTCCTCCGAAAGTGAGTATGAAGAAGTGGATATCAATGATATATATGTACCAAGTTTTCCCAAATATAAAACGTTCATTGAATTAGTACTAGAACCTTCCAAAAGGGATACATTTAATACATCAAGTGGTGACACATTCACCAATAAACTTACGGATGATGAATGGAACCAATTGAAACAGGATTTTATTGAACAATATTTACAAAACATACAAAAGGATTTTATTTTACATGATAGTATGGATGAAAAACCTTTTATTACTCAAATCCAGGATAGATTTCTTGATAGTAGTCATGAAGAAGTTACTTATAATATTGATTGGAATGTTCCTGAAAATATTAATAGGATTACTAATAACATGGACGATCCAAAATACTGCTCAAATAATATGTATACTGGTACCGATTTAATTAATGATTCATTAAATGGTAACCAATATATTGATATATATGATGAGATGCTGAAACGAAAAGAAAACGAATTATTTGGAACATATCATACAAAATATACAACCTTTAACAGTGTTTCTAAACAAACACCTAGTGACCCGATAATTAACCAACTAGATTTATATCATAAATGGATAGACAAACATAGAGATATTTGCGAACAGTGGAAAACCAAAGAGGATATGTTATATAAATTGAATGAAGTGTGGAATATGGAACGTAAGGAATATCTATTGGATATACAACCATCAACTCTGGATGATATTCATAAAATTAATGATGAAACATATAATATTATTAGTACAAATAATATATATGATCATCCCTCACAGGAAACCCCCCTCCAACTACTTGGATCAACAAATATTATACCCAGTTATATTACCACGGAACAAAATAATGGATTGCGCACAAATATATCTATGGATACATATATTGATGAAACAAATAATAATAATGTGGTAGCCACTAGTATAATAGGTGACGATCAGATGGAAAATTCGTACAATTGTCCCGGGATGGTAAGGCCTTTGAATTGTATAGTTGCAGTTAGTCAAAATATGGGAATTGGGAAAAATGGTGATCTTCCATGGCCCCCCTTACGTAATGAGTTCAAGTACTTTCAAAGGATGACTACCACAAGTAGTGTGGAGGGAAAGCAAAACTTAGTAATAATGGGGCGTAAAACTTGGTTTAGTATACCCGAAAAAAATAGGCCATTGAAAGATCGTATAAACATTGTATTGTCACGTGAGTTGAAAGAGCCACCTAGAGGAGCACACTTCTTGGCTAAATCTTTGGACGACGCTTTGAGGTTAATAGAGCAGCCTGAGCTTGCTTCAAAGGTTGATATGGTATGGATTGTAGGAGGTTCAAGTGTATACCAAGAGGCAATGAACCAACCTGGACACTTAAGGTTGTTCGTAACTCGTATTATGCAGGAGTTCGAGTCAGATACATTCTTCCCTGAGATAGACCTTGGAAAGTACAAGTTGTTACCTGAGTATCCTGGAGTATTAAGTGAAGTTCAAGAAGAAAAGGGAATAAAATATAAGTTCGAGGTTTACGAAAAAAAGGAC

>3D7var0425800

TACCACGAATACATTTACAGATGAGGAATGGAATGAACTGAAACAGGATTTTGTATCACAATATATACAAAGTAGATTACCAATGGATGTACCACAATATGATGTATCAACGGAGAGTCCAATGAATATAGGAGGTAATGTTTTAGATGATGGTATGGATGAAAAACCTTTTATTACTTCTATTCATGATAGGGATTTAAATAGTGGAGAAGAAATTAGTTATAATATTCATATGAGTACTAACACTAATAATGATATTCCAAAATATGTATCAAATAATGTATATTCTGGTATAGATTTAATTAATGATACATTAAGTGATAACAAACATATTGATATATATGATGAAGTGCTAAAAAGAAAAGAAAATGAATTATTTGGAACAAATTATAAGAAAAATACATCAAACAATAGTGTAGCAAAAAATACTAATAGTGATCCAATTATGAACCAATTAGATTTGTTACATAAATGGTTAGATAGACATAGAGATATATGTGAAAATTGGGGGAAAAAAGAAGATATTTTGAATAAATTGAATGAACAATGGAATAAAGATAATGATGGTGGTGATATACCAAATGATAACAAAAAGTTGAATACGGATGTTTCGATACAAATAGATATGGATGAAACTAAAGGAAAGAAGGAATTTAGTAATATGGATACTATCTTGGATGATATGGAAGATGATATATATTATGATGTAAATGATGAAAACCCATCTGTAGATGATATACCTATGGATCATAATAAAGTAGATGTACCAAAGAAAGTACATGTTGAAATGAAAATCCTTAATAATACATCTAATGGATCGTTGGAACAACAATTTCCTATATCGGATGTATGGAATATA

>3D7rif0425700

GGATGTACCACAATATGATGTATCAACGGAGAGTCCAATGAATATAGGAGGTAATGTTTTAGATGATGGTATGGATGAAAAACCTTTTATTACTTCTATTCATGATAGGGATTTAAATAGTGGAGAAGAAATTAGTTATAATATTCATATGAGTACTAACACTAATAATGATATTCCAAAATATGTATCAAATAATGTATATTCTGGTATAGATTTAATTAATGATACATTAAGTGATAACAAACATATTGATATATATGATGAAGTGCTAAAAAGAAAAGAAAATGAATTATTTGGAACAAATTATAAGAAAAATACATCAAACAATAGTGTAGCAAAAAATACTAATAGTGATCCAATTATGAACCAATTAGATTTGTTACATAAATGGTTAGATAGACATAGAGATATATGTGAAAATTGGGGGAAAAAAGAAGATATTTTGAATAAATTGAATGAACAATGGAATAAAGATAATGATGGTGGTGATATACCAAATGATAACAAAAAGTTGAATACGGATGTTTCGATACAAATAGATATGGATGAAACTAAAGGAAAGAAGGAATTTAGTAATATGGATACTATCTTGGATGATATGGAAGATGATATATATTATGATGTAAATGATGAAAACCCATCTGTAGATGATATACCTATGGATCATAATAAAGTAGATGTACCAAAGAAAGTACATGTTGAAATGAAAATCCTTAATAATACATCTAATGGATCGTTGGAACAACAATTTCCTATATCGGATGTATGGAATATA

>3D7rif1254800

GGATGTACCACAATATGATGTATCAACGGAGAGTCCAATGAATATAGGAGGTAATGTTTTAGATGATGGTATGGATGAAAAACCTTTTATTACTTCTATTCATGATAGGGATTTAAATAGTGGAGAAGAAATTAGTTATAATATTCATATGAGTACTAACACTAATAATGATATTCCAAAATATGTATCAAATAATGTATATTCTGGTATAGATTTAATTAATGATACATTAAGTGATAACAAACATATTGATATATATGATGAAGTGCTAAAAAGAAAAGAAAATGAATTATTTGGAACAAATTATAAGAAAAATACATCAAACAATAGTGTAGCAAAAAATACTAATAGTGATCCAATTATGAACCAATTAGATTTGTTACATAAATGGTTAGATAGACATAGAGATATATGTGAAAATTGGGGGAAAAAAGAAGATATTTTGAATAAATTGAATGAACAATGGAATAAAGATAATGATGGTGGTGATATACCAAATGATAACAAAAAGTTGAATACGGATGTTTCGATACAAATAGATATGGATGAAACTAAAGGAAAGAAGGAATTTAGTAATATGGATACTATCTTGGATGATATGGAAGATGATATATATTATGATGTAAATGATGAAAACCCATCTGTAGATGATATACCTATGGATCATAATAAAGTAGATGTACCAAAGAAAGTACATGTTGAAATGAAAATCCTTAATAATACATCTAATGGATCGTTGGAACAACAATTTCCTATATCGGATGTATGGAATATA

>IT4var66

CACATTAAGTGGTAATGAACATATTGATATATATGATGAATTGTTGAAACGAAAAGAAAACGAATTGTTCGGAACTAATCATCCAAAACGTACATCAAACAATAGTGTTATTAAATCAACAAATAGTGATCCTATACTCAATCAAATAAATTTGTTCCATACATGGCTAGATAGACATAAAAATATGTGCGAACAGTGGGATAAAAATAAAAAGGAGGAATTGTTAGATAAATTGAAAGAACAGTGGGAAAATGAGACACATAGTGGTAACACTCACCCTAGTGATAGTAACAAAACGTTGAATACTGATGTTTCTATTCAAATACATATGGATAATCCTAAACCTATAAATCAATTTACTAATATGGATACAAGCCCCGACAAATCTACTATGGATACAATAATTGATGATCTGGAAAAATATAACGAACCCTACTACTATTATTTTTATAAAGATGATATCTATTATGATGTAAATGATGATGATAAAACATCTATGGACAACAACAATAACCTTGTAAATAAGAATAACCCTGTAGATAGTAACAGTTCCACCTATAACCACCACAACCCTGCAGACATTAACAAAACTTTTGTGGATATAAACAATCATAACCAACACCCTATAGAAAAACCTACCAAAATACAAATTGAAATGAATTCAAATAATCGCGAAGTGGACGAACAGCAATATCCTATAGCGGATATATGGAATATA

>IT4var2csa

TAAATATATTTGCGACTTATCTTCCTCTGATATTACATCTTCGTCAGAGAGCGAATATGAAGAATTGGATATCAATGATATATATGTACCAAGTCTTCCCAAATATAAAACGTTGATTGAATTAGTACTAGAACCTTCAAAAAGGGATACATTTAATACACCAAGTGGTGACACATTCACCAATAAATTTAGAGATGATGAATGGAACCAACTGAAACAGGATTTTATTGAACAATATTTACAAAACATACAAAAGGATTTTATTTTACATTATAGTATGGATGAAAAACCTTTTATTACTCAAATACAAGATAGATTTCTGGATAGTAGTCATGAAGAAGTTATTTATAATATTGATTGGAATGTTCCTGAAAATATTAATAGGATTACTAATATCATGGACGATCCAAAATACTCCTCAAATAATATGTATACTGGTACCGATTTAATTAATGATTCATTAAATGGTAACCAACATATGGATATATATGATGAAATGCTCAAACGAAAAGAAAATGAATTATTTGGAACAAATAATACAAAAAATACAACATTTAATAGTGTTTATAAACAAACACATAGTGACCCGATAATTAACCAACTAAATTTATATCATAAATGGTTAGACAAACATAGAGATATTTGCGAACAGTGTAAAACGAAAAAGGATATGTTATATAAATTGAATGAAGTGTGGAATATGGAACGTAAGGAATATCTATTGGATATACCTCCATCAATTCTGGATGATATTCATAAAACTAATGATGAAACATATAATATTATTAGTACAAATAATATATATGATCATCCCTCACAGGAAACCCCCCTCCAACTACTTGGATCAACAAATATTATACCCAGTTATATTACCACGGAACAAAATAATGGATTGCGAACAAATATATCTATGGATATACATATTGATGAAAAAAATTATAATAATGTTGTAGCCACTAGTATAATAGGTGACGATCAGGTGGAAAATTCGTACAATTTG

>IT4var01

TTGGAGACATATCTTCATCTGATATTACTTCATCAGAAAGTGAGTATGAAGAATTGGATATCAATGATATATACCCATACAAATCACCTAAATATAAAACGTTGATTGAAGTGGTACTAGAACCATCAAAAAGAGATACAATGAACACGCAAAGTGATATACCATTAAATGATAAACTTGATAGTAATAAACTTACAGATGAAGAATGGAATCAACTGAAACAGGATTTTATTTCAAATATTTCACAAAATTCTCAAATGGATTTACCCAAAAATAATATAAGTGGGAATATTCAAATGGATACCCATCCTCATGTTAATATTTTAGACGATAGTATGCAAGAAAAACCTTTTATTACATCTATTCATGATAGAGATTTACATAATGGTGAAGAAGTTACCTATAATATTAATTTGGATGATCACAAAAATATGAATTTTTCAACTAATCATGATAATATACCACCAAAAAATGATCAAAATGATTTATATACTGGTATAGATTTGATTAATGATTCGATAAGTGGTAACCATAATGTTAATATTTATGATGAATTGTTAAAAAGAAAAGAAAACGAATTATTTGGAACAAATCATACAAAACATACAACAACAAATATTGTTGCCAAACAAACACATAATGACCCTATAGTCAATCAAATAAATTTGTTCCATAAATGGTTAGATAGACATAGAAATATGTGCGAACAGTGGGATAAAAATAAAAAGGAGGAATTGTTAGATAAATTGAATGAAGAATGGAATAAAGAAAATAAAAATAATAGTAATGTCACAGACACAAATGGTGAAAATAATATTACAAGGGTGTTGAATAGTGATGTTTCTATCCAAATAGATATGAATTCTAAACCTATT

>IT4var16

TAGTGGTAAGAACACAACAGCTAGTGGTAAAAACACACCTAGTGATACACAAAATGATATACAAAATGATGGTATACCTAGTAGTGATACACCTATGAATAAATTTAATGATGATGAATGGAATCAATTGAAACATGATTTTATATCTAATATGTTACAAAATACCCAAAATAAGGAACCAAATATTTTACATGATAATGTGGATAATAATACCCATCCTACCATGTCACGTCATAATATAGACCAAAAACCTTTTATTATGTCCATACATGATAGAAATTTATATATTGGACAAGAATATAGTTATGATATGAGTACTAACAGTGCTAATAATGATTTATATAGTGGTCAAAATAATTTATATAGTGATGTAGATTCAACAAGTGGAAACCGTGATTCATATAGTGATAAAAATGATCCAATTAGTGATAACCATCATCCTTATAGTGGTATTGACCTAATTAATGATTCATTAAATAGTGGTAATCATGACATATATGATGAAATATTAAAACGAAAAGAAAACGAATTATTTGGGACCAAACATCACCCAAAACATACAAATATATATAATGTCGCCAAACCTGCACGTGACGACCCTATAACCAATCAAATAAATTTGTTCCATAAATGGTTAGATAGACATAGAGACATGTGCGAGAAGTGGGATACAAACAATAAAGTGGATATTTTGAACCAATTGAAAGAAGAGTGGGAAAATGATAATAGCAATAGTGGTAATAAAACTAGTGGAAATATCACACCAACTAGTGATATACCTAGTGGTAAACTAAGTGATATACCTAGTACTAACAAAATGTTGAATAGTGATGTTTCTATTCAAATACATATAGATAAGCCCAACCAAGTGGATGATAACATATACCTTGATACATACCCCGATAAATATACTGTGGATAACATTAACCCTGTCGATACACACACCAACCCAAACCTTGTGGGAAACATCAACCCTGTGGATCAAAACTCCAACCTAACGTTTCCGTCTAATCCAAACCCTGCGTATGATAACATATATATTGATCATAATAATGAGGATCTACCTAGCAAAGTACAAATTGAAATGAGTGTGAAGAACGGAGAAATGGCGAAAGAGAAA

>IT4var19

CTATGAATAAATTTACTGATGAGGAATGGAATCAATTGAAACACGATTTTATATCTCAATATTTACCAAATACAGAACCTAATACTTTATATTTTGATAAACCTGAAGAAAAACCTTTTATTACTTCTATTCATGATAGAAATTTATATACTGGGGAAGAATATAGTTATAATATTAATATGAGTACTAATACTATGGATGATCCAAAATATGTATCAAATAATGTATATTCTGGTATTGACCTAATTAATGATTCATTAAATAGTGGTAATCAACCTATTGATATATATGATGAAGTGTTGAAACGAAAAGAAAATGAATTATTCGGGACAGAACATCACCCAAAACGTACAACAACTAACCATTTCGCCACACCTACACGTGACGACCCCATCCACAACCAACTGGAACTATTCCATAAATGGTTAGATCGACATAGAGATATGTGCGAGAAGTGGAATAATAAAGAGGAAGTATTAGATAAATTAAAAGAAGAGTGGGAAAATGAGACACATAGTGGTAACACTCACCCTAGTGATAGTAACAAAACGTTGAATACGAATGTTTCTATTCAGATAGATATGGATCATGAAAAACGAATGAAGGAATTTACTAATATGGATACATATCCGGAGAATTCTACTATGGATAGTATATTGGAGGATCTGGAAAAATATAAAGAACCTTATTATGATGTGCAAGATGATATTTATTATGATGTAAATGATCATGATACATCAACTGTGGATAGTAATAATATGGATGTTCCTAGTAAAGTACAAATTGAAATGGATGTAAATACCAAATTGGTTAAAGAGAAATATCCTATATCGGATGTATGGGATATA

>IT4var01BirA*Pos1

TTGGAGACATATCTTCATCTGATATTACTTCATCAGAAAGTGAGTATGAAGAATTGGATATCAATGATATATACCCATACAAATCACCTAAATATAAAACGTTGATTGAAGTGGTACTAGAACCATCAAAAAGAGATACAATGAACACGCAAAGTGATATACCATTAAATGATAAACTTGATAGTAATAAACTTACAGATGAAGAATGGAATCAACTGAAACAGGATTTTATTTCAAATATTTCACAAAATTCTCAAATGGATTTACCCAAAAATAATATAAGTGGGAATATTCAAATGGATACCCATCCTCATGTTAATATTTTAGACGATAGTATGCAAGAAAAACCTTTTATTACATCTATTCATGATAGAGATTTACATAATGGTGAAGAAGTTACCTATAATATTAATTTGGATGATCACAAAAATATGAATTTTTCAACTAATCATGATAATATACCACCAAAAAATGATCAAAATGATTTATATACTGGTATAGATTTGATTAATGATTCGATAAGTGGTAACCATAATGTTAATATTTATGATGAATTGTTAAAAAGAAAAGAAAACGAATTATTTGGAACAAATCATACAAAACATACAACAACAAATATTGTTGCCAAACAAACACATAATGACCCTATAGTCAATCAAATAAATTTGTTCCATAAATGGTTAGATAGACATAGAAATATGTGCGAACAGTGGGATAAAAATAAAAAGGAGGAATTGTTAGATAAATTGAATGAAGAATGGAATAAAGAAAATAAAAATAATAGTAATGTCACAGACACAAATGGTGAAAATAATATTACAAGGGTGTTGAATAGTGATGTTTCTATCCAAATAGATATGAATTCTAAACCTATTGGTACCGGAGGTGGAGGTAGTGGAGGCGGAGGTAGCGGAGGCGGTGGGAGTGGTGGAGGCGGAAGTGGCGGTGGAGGTAGCGGAGGTGGAGGTAGTGGAGGCGGAGGTAGCATGAAAGATAATACAGTACCATTAAAATTAATAGCTTTATTAGCTAATGGTGAATTTCATTCAGGTGAACAATTAGGTGAAACATTAGGTATGTCAAGAGCTGCTATAAATAAACATATACAAACATTAAGAGATTGGGGTGTAGATGTATTTACAGTACCAGGTAAAGGTTATTCATTACCAGAACCAATACAATTATTAAATGCTGAAAAAATATTATCACAATTAGATGATGGTTCAGTAGCTGTATTACCAGTAATAGATTCAACAAATCAATATTTATTAGATAGAATAGGTGAATTAAAATCAGGTGATGCTTGTGTAGCTGAATATCAACATGCTGGTAGAGGTGGTAGAGGTAGAAAATGGTTTTCACCATTTGGTGCTAATTTATATTTATCAATGTTTTGGAGATTAGAACAAGGTCCAGCTGCTGCTATAGGTTTATCATTAGTAATAGGTATAGTAATGGCTGAAGTATTAAGAAAATTAGGTGCTGATAAAGTAAGAGTAAAATGGCCAAATGATTTATATTTACAAGATAGAAAATTAGCTGGTATATTAGTAGAATTAACAGGTAAAACAGGTGATGCTGCTCAAATAGTAATAGGTGCTGGTATAAATATGGCTATGAGAAGAGTAGAAGAATCAGTAGTAAATCAAGGTTGGATAACATTACAAGAAGCTGGTATAAATTTAGATAGAAATACATTAGCTGCTATGTTAATAAGAGAATTAAGAGCTGCTTTAGAATTATTTGAACAAGAAGGTTTAGCTCCATATTTATCAAGATGGGAAAAATTAGATAATTTTATAAATAGACCAGTAAAATTAATAATAGGTGATAAAGAAATATTTGGTATATCAAGAGGTATAGATAAACAAGGTGCTTTATTATTAGAACAAGATGGTATAATAAAACCATGGATGGGTGGTGAAATATCATTAAGATCAGCTGAAAAA

>IT4var01BirA*Pos2

TGTATACAGAATAGTGGAAATGAAAAAAAGTGGGAACATGCTTTAGATACGATAAAAATAAAAAATGGTGCACCTACTAGTATTAATGTCCAAATGATTGATCGTAGGGGACAGTACATTCAAGAGCATTCAGAAAATTCTTTTAAAGAATCACGTCTTTTAAAAAGTGTCAGAGAACAAAAATGGGAATGTAGCTTTGTTAATAAAAAGATGGATGTATGTAAACTAAAGAATTTTAAGGAAAACATAGACACTGATGAAACCATTACATTTAAAGTACTCCTAGAGAATTGGTTACAAGATTTCATAGAAGGTTATTATATATCAAAAAGGAAAATCGATATATGTACAAAAAAAGAAGAACATACAGCAATTGAAGGATGTAAAAGTAAATGCGAATGTATTGGAAAATGGTTAAAGCAAAAGACTACAGAATGGGATGAAATAAAAACACATTTTAATAAACAAAACCGCGGTGATGGATATGAAATAGCTCATAAGGTCAGAAATTATTTTGAGAAAAATGCAGTTCAATTAAAAAAATGGATAGATGATCTTAAACATGTAAAAAAAATAGATGATTCTGAGGATTGTAGTGTTAATAACGATTGCGCAAATATGGATAAAGAAACCAATAAAGATATGGTATCTATTTTACTTTCTCAGCTTAAAAAAGAAATAAAACCTTTCGAAAATCAACCTCACGAAACGACTTCTCCAAATTATTGTGATATATCCCCCACACACATACTCACAGAACCCTCCCAAACTGATGACACCGAAACCCTTAACCCTTACGACGAAACCCCAGAAGACGACACATCCACATCTTCACGTCCGAATTTTTGTCCGCAAGTGGAACCACCACCCAAACCGGAAGTGCCACAAAAACCTGAAGAAACTGCAGAAGATACCACAGAAGACACGGAGGAAGAAGCTGCAGCTCCACCCGTAGCTCCATCTAGTGGCGAAGAAGAAGCTCCTAAAGAAGTAGTACCAGAGAAAAAACCAAAAGAAGTACCAAAACCAGGACCAAAAGCACCAAAAAAACGACAACCACGTGAAGTGACGCATTCCATAGTCACTAACCTGTTACTACCATCGGCCTTCCCGCTAAGTGTAGGAATCGCGTTTGCTGCGTTGAGTTATTTCGTACTAAAGgtaagaactatgtgcatatgtgtatatgtgtgtgttgtatgtgtggatatatacgtatgcgttttatatatattttatatatatttatattaaaaaaggaaaaaagaaaaaggaatatagaaacataattgttaaaaaaaaaattaaaaaaatgttagaaaaaaaattaaaaaaatgtaaaaaaaaaaaaataaaaaaaaaaaaataaataaataaattaaaaaaaaataataataaattacatacatacacatatacatacatatattatatacataccaaaacctacatacatatatgttcgttttttttagAAAAAAACCAAAGGTACCGGTGGAGGTGGATCAGGTGGAGGTGGATCAATGAAAGATAATACAGTACCATTAAAATTAATAGCTTTATTAGCTAATGGTGAATTTCATTCAGGTGAACAATTAGGTGAAACATTAGGTATGTCAAGAGCTGCTATAAATAAACATATACAAACATTAAGAGATTGGGGTGTAGATGTATTTACAGTACCAGGTAAAGGTTATTCATTACCAGAACCAATACAATTATTAAATGCTGAAAAAATATTATCACAATTAGATGATGGTTCAGTAGCTGTATTACCAGTAATAGATTCAACAAATCAATATTTATTAGATAGAATAGGTGAATTAAAATCAGGTGATGCTTGTGTAGCTGAATATCAACATGCTGGTAGAGGTGGTAGAGGTAGAAAATGGTTTTCACCATTTGGTGCTAATTTATATTTATCAATGTTTTGGAGATTAGAACAAGGTCCAGCTGCTGCTATAGGTTTATCATTAGTAATAGGTATAGTAATGGCTGAAGTATTAAGAAAATTAGGTGCTGATAAAGTAAGAGTAAAATGGCCAAATGATTTATATTTACAAGATAGAAAATTAGCTGGTATATTAGTAGAATTAACAGGTAAAACAGGTGATGCTGCTCAAATAGTAATAGGTGCTGGTATAAATATGGCTATGAGAAGAGTAGAAGAATCAGTAGTAAATCAAGGTTGGATAACATTACAAGAAGCTGGTATAAATTTAGATAGAAATACATTAGCTGCTATGTTAATAAGAGAATTAAGAGCTGCTTTAGAATTATTTGAACAAGAAGGTTTAGCTCCATATTTATCAAGATGGGAAAAATTAGATAATTTTATAAATAGACCAGTAAAATTAATAATAGGTGATAAAGAAATATTTGGTATATCAAGAGGTATAGATAAACAAGGTGCTTTATTATTAGAACAAGATGGTATAATAAAACCATGGATGGGTGGTGAAATATCATTAAGATCAGCTGAAAAAGGAAGTGGAAGTGGAAGTACGCGTAGCACAATAGACTTACTTAGAGTAATAGACATTCCAAAGGGTGACTACGGTATTCCAACTCTTAAGAGTAGTAACCGTTACATACCATACGGAACAAACAGGTACAAGGGTAAGACTTATATATACATGGAGGGTGACTCAGGTGATGAGAAGTACATAGGTGATATTAGCAGTAGCGACATAACAAGTAGTGAGTCAGAATACGAGGAGCTTGACATAAACGACATTTATCCTTATAAGAGTCCAAAGTACAAGACACTTATAGAGGTTGTTTTGGAGCCTAGTAAGCGTGACACTATGAATACACAGTCAGACATTCCTCTTAACGACAAGTTAGACTCAAACAAGTTAACTGACGAGGAGTGGAACCAGTTAAAGCAAGACTTCATAAGTAACATAAGTCAGAACAGCCAGATGGACCTTCCAAAGAACAACATTTCAGGAAACATACAGATGGACACACACCCACACGTAAACATACTTGATGACTCAATGCAGGAGAAGCCATTCATAACTAGCATACACGACCGTGACCTTCACAACGGAGAGGAGGTAACATACAACATAAACCTTGACGACCATAAGAACATGAACTTCAGTACAAACCACGACAACATTCCTCCTAAGAACGACCAGAACGACCTTTACACAGGAATTGACCTTATAAACGACAGTATTTCAGGAAATCACAACGTAAACATATACGACGAGCTTCTTAAGCGTAAGGAGAATGAGCTTTTCGGTACTAACCACACTAAGCACACTACTACTAACATAGTAGCAAAGCAGACTCACAACGATCCAATTGTTAACCAGATTAACCTTTTTCACAAGTGGCTTGACCGTCACCGTAACATGTGTGAGCAATGGGACAAGAACAAGAAAGAAGAGCTTCTTGACAAGCTTAACGAGGAGTGGAACAAGGAGAACAAGAACAACTCAAACGTTACTGATACTAACGGAGAGAACAACATAACTCGTGTTCTTAACTCAGACGTAAGCATACAGATTGACATGAACAGCAAGCCAATA

>IT4var01BirA*Pos3

GGAACATCTAAAAACATCATCTAATCTTGATAATAAATATGTTAAAGAATTTTATGCAACATCTGAAGGAAAATATAAATCTGTTGACTCATTTTTAGATAAATTGAAAGAAAGATCCCATTGTCATATGGATACGCTAGAAGGAAAAATAGATTTTAAGAATCCGCTCAAAACGTTTTCTTCTTCAACATATTGTAAAACGTGCCCTTTATATGGTGTTCAGTGTAGGAATACATCTGATCACTGTATACAGAATAGTGGAAATGAAAAAAAGTGGGAACATGCTTTAGATACGATAAAAATAAAAAATGGTGCACCTACTAGTATTAATGTCCAAATGATTGATCGTAGGGGACAGTACATTCAAGAGCATTCAGAAAATTCTTTTAAAGAATCACGTCTTTTAAAAAGTGTCAGAGAACAAAAATGGGAATGTAGCTTTGTTAATAAAAAGATGGATGTATGTAAACTAAAGAATTTTAAGGAAAACATAGACACTGATGAAACCATTACATTTAAAGTACTCCTAGAGAATTGGTTACAAGATTTCATAGAAGGTTATTATATATCAAAAAGGAAAATCGATATATGTACAAAAAAAGAAGAACATACAGCAATTGAAGGATGTAAAAGTAAATGCGAATGTATTGGAAAATGGTTAAAGCAAAAGACTACAGAATGGGATGAAATAAAAACACATTTTAATAAACAAAACCGCGGTGATGGATATGAAATAGCTCATAAGGTCAGAAATTATTTTGAGAAAAATGCAGTTCAATTAAAAAAATGGATAGATGATCTTAAACATGTAAAAAAAATAGATGATTCTGAGGATTGTAGTGTTAATAACGATTGCGCAAATATGGATAAAGAAACCAATAAAGATATGGTATCTATTTTACTTTCTCAGCTTAAAAAAGAAATAAAACCTTTCGAAAATCAACCTCACGAAACGACTTCTCCAAATTATTGTGATATATCCCCCACACACATACTCACAGAACCCTCCCAAACTGATGACACCGAAACCCTTAACCCTTACGACGAAACCCCAGAAGACGACACATCCACATCTTCACGTCCGAATTTTTGTCCGCAAGTGGAACCACCACCCAAACCGGAAGTGCCACAAAAACCTGAAGAAACTGCAGAAGATACCACAGAAGACACGGAGGAAGAAGCTGCAGCTCCACCCGTAGCTCCATCTAGTGGCGAAGAAGAAGCTCCTAAAGAAGTAGTACCAGAGAAAAAACCAAAAGAAGTACCAAAACCAGGACCAAAAGCACCAAAAAAACGAGGTACCGGAAGTGGAAGTGGAAGTATGAAAGATAATACAGTACCATTAAAATTAATAGCTTTATTAGCTAATGGTGAATTTCATTCAGGTGAACAATTAGGTGAAACATTAGGTATGTCAAGAGCTGCTATAAATAAACATATACAAACATTAAGAGATTGGGGTGTAGATGTATTTACAGTACCAGGTAAAGGTTATTCATTACCAGAACCAATACAATTATTAAATGCTGAAAAAATATTATCACAATTAGATGATGGTTCAGTAGCTGTATTACCAGTAATAGATTCAACAAATCAATATTTATTAGATAGAATAGGTGAATTAAAATCAGGTGATGCTTGTGTAGCTGAATATCAACATGCTGGTAGAGGTGGTAGAGGTAGAAAATGGTTTTCACCATTTGGTGCTAATTTATATTTATCAATGTTTTGGAGATTAGAACAAGGTCCAGCTGCTGCTATAGGTTTATCATTAGTAATAGGTATAGTAATGGCTGAAGTATTAAGAAAATTAGGTGCTGATAAAGTAAGAGTAAAATGGCCAAATGATTTATATTTACAAGATAGAAAATTAGCTGGTATATTAGTAGAATTAACAGGTAAAACAGGTGATGCTGCTCAAATAGTAATAGGTGCTGGTATAAATATGGCTATGAGAAGAGTAGAAGAATCAGTAGTAAATCAAGGTTGGATAACATTACAAGAAGCTGGTATAAATTTAGATAGAAATACATTAGCTGCTATGTTAATAAGAGAATTAAGAGCTGCTTTAGAATTATTTGAACAAGAAGGTTTAGCTCCATATTTATCAAGATGGGAAAAATTAGATAATTTTATAAATAGACCAGTAAAATTAATAATAGGTGATAAAGAAATATTTGGTATATCAAGAGGTATAGATAAACAAGGTGCTTTATTATTAGAACAAGATGGTATAATAAAACCATGGATGGGTGGTGAAATATCATTAAGATCAGCTGAAAAAGGTGGAGGTGGATCAGGTGGAGGTGGATCAACCGGTCAGCCTAGAGAGGTTACACACAGTATTGTTACAAATTTACTTTTGCCTAGTGCATTTCCATTGTCAGTTGGTATAGCATTCGCAGCACTTTCATACTTTGTTTTGAAAAAGAAGACAAAGAGCACAATAGACTTACTTAGAGTAATAGACATTCCAAAGGGTGACTACGGTATTCCAACTCTTAAGAGTAGTAACCGTTACATACCATACGGAACAAACAGGTACAAGGGTAAGACTTATATATACATGGAGGGTGACTCAGGTGATGAGAAGTACATAGGTGATATTAGCAGTAGCGACATAACAAGTAGTGAGTCAGAATACGAGGAGCTTGACATAAACGACATTTATCCTTATAAGAGTCCAAAGTACAAGACACTTATAGAGGTTGTTTTGGAGCCTAGTAAGCGTGACACTATGAATACACAGTCAGACATTCCTCTTAACGACAAGTTAGACTCAAACAAGTTAACTGACGAGGAGTGGAACCAGTTAAAGCAAGACTTCATAAGTAACATAAGTCAGAACAGCCAGATGGACCTTCCAAAGAACAACATTTCAGGAAACATACAGATGGACACACACCCACACGTAAACATACTTGATGACTCAATGCAGGAGAAGCCATTCATAACTAGCATACACGACCGTGACCTTCACAACGGAGAGGAGGTAACATACAACATAAACCTTGACGACCATAAGAACATGAACTTCAGTACAAACCACGACAACATTCCTCCTAAGAACGACCAGAACGACCTTTACACAGGAATTGACCTTATAAACGACAGTATTTCAGGAAATCACAACGTAAACATATACGACGAGCTTCTTAAGCGTAAGGAGAATGAGCTTTTCGGTACTAACCACACTAAGCACACTACTACTAACATAGTAGCAAAGCAGACTCACAACGATCCAATTGTTAACCAGATTAACCTTTTTCACAAGTGGCTTGACCGTCACCGTAACATGTGTGAGCAATGGGACAAGAACAAGAAAGAAGAGCTTCTTGACAAGCTTAACGAGGAGTGGAACAAGGAGAACAAGAACAACTCAAACGTTACTGATACTAACGGAGAGAACAACATAACTCGTGTTCTTAACTCAGACGTAAGCATACAGATTGACATGAACAGCAAGCCAATA

SLI2 integration with Ty1 and TGD constructs

xxxxx = mScarlet

xxxxx = Skip peptide

xxxxx = yDHODH

xxxxx = BD-Resistance

>pSLI2a-**Insert**-mScarlet-T2A-yDHODH

**gcggccgcINSERTGTCGACGGAGAAGGAAGAGGAAGTTTATTAAC**ATGTGGAGATGTAGAAGAAAATCCAGGACCAATGACAGCCAGTTTAACTACCAAGTTCTTGAACAATACCTATGAAAACCCATTTATGAATGCATCCGGTGTTCATTGCATGACTACACAAGAATTAGATGAATTAGCAAACTCTAAAGCTGGCGCATTCATTACAAAGAGTGCTACAACCTTAGAAAGAGAAGGTAACCCTGAACCACGTTACATTTCTGTCCCTCTAGGCAGTATCAACTCCATGGGTTTACCAAACGAAGGTATCGACTACTATTTGTCCTATGTATTAAACCGTCAAAAGAATTATCCTGATGCACCTGCTATTTTCTTCTCAGTTGCTGGTATGAGCATTGATGAAAATTTAAATTTGTTGAGGAAAATCCAAGATAGCGAATTCAACGGTATTACCGAGTTAAACTTGTCTTGTCCTAATGTGCCTGGGAAACCACAAGTTGCTTATGACTTTGACTTGACAAAGGAAACCTTGGAAAAGGTTTTTGCCTTTTTCAAAAAACCTCTTGGTGTCAAGTTGCCTCCTTATTTTGATTTTGCCCATTTTGATATCATGGCAAAAATATTGAACGAGTTCCCATTAGCTTATGTCAACTCTATCAATAGTATAGGAAATGGTCTTTTCATTGATGTGGAGAAGGAGAGTGTAGTAGTGAAGCCAAAGAATGGTTTCGGGGGTATTGGAGGTGAATATGTTAAGCCAACCGCGCTCGCCAATGTTCGTGCATTTTACACTCGTTTGAGACCTGAAATCAAAGTTATCGGTACAGGTGGAATTAAGTCCGGTAAGGATGCATTTGAACATCTTCTATGTGGTGCCTCTATGCTACAGATTGGTACAGAATTACAAAAAGAGGGCGTCAAGATTTTTGAACGTATCGAAAAAGAATTAAAAGACATAATGGAAGCTAAGGGTTATACATCCATAGATCAGTTCCGTGGGAAGTTGAACAGCATTTAACCCGGGtcgagggatatggcagcttaatgttcgtttttcttatttatatatttataccaattgattgtatttataactgtaaaaatgtgtatgttgtgtgcatatttttttttgtgcatgcacatgcatgtaaatagctaaaattatgaacattttattttttgttcagaaaaaaaaaactttacacacataaaatggctagtatgaatagccatattttatataaattaaatcctatgaatttatgaccatattaaaaatttagatatttatggaacataatatgtttgaaacaataagacaaaattattattattattattatttttactgttataattatgtgtctccttcaatgattcataaatagttggacttgatttttaaaatgtttataatatgattagcatagttaaataaaaaaagttgaaaaattaaaaaaaaacatataaacacaaatgatggtttttccttcaatttcgatatcaatttatagaaacaaaatatatacttgtataattttatttttttatataaatcattacatatataattatacaatattttttctaagagataattatatattaatatatataaaaaaaggtgttttttttttttttttttatttttatttttattttatggtaatattttattttccttattttataaattatattagtttatatgtgattaattttatatattatcaatttatatatttttaaatgcttacttaattatctttttttttttttttttttttttttcccctctttttatattaatttatttttgaaaaaattgatatatatatatatatataatatatatatatacatgtagtagtattaaacaatgtataatatatataaataatatatttatatatttcatttcaattttaattttttttggttttttttttttttctttttgtcatatttaaaaaaaattatattcatataagttatgcattttttataaacattattcaatatatgtataatataatatatatatatatattaatgtattattccaatgtgcatgataaaagaaaaaaataatatttataaaaaaaaagaaaaataaaacaaaaaaagaaaaaaaaaaaaaaaaaaaaaaaaatacaaaaataaataatataatttataattatatattcttgtcacaataaaaatatatatatatatatatatatttataatatgtatattttaaactagaaaaggaataactaatattttatttattatcattcaagatttatattttataataataaatacctaatagaaatatatcaGGATCCATGGCCAAGCCTTTGTCTCAAGAAGAATCCACCCTCATTGAAAGAGCAACGGCTACAATCAACAGCATCCCCATCTCTGAAGACTACAGCGTCGCCAGCGCAGCTCTCTCTAGCGACGGCCGCATCTTCACTGGTGTCAATGTATATCATTTTACTGGGGGACCTTGTGCAGAACTCGTGGTGCTGGGCACTGCTGCTGCTGCGGCAGCTGGCAACCTGACTTGTATCGTCGCGATCGGAAATGAGAACAGGGGCATCTTGAGCCCCTGCGGACGGTGCCGACAGGTGCTTCTCGATCTGCATCCTGGGATCAAAGCGATAGTGAAGGACAGTGATGGACAGCCGACGGCAGTTGGGATTCGTGAATTGCTGCCCTCTGGTTATGTGTGGGAGGGCAAGCTTatttaataatagattaaaaatattataaaaataaaaacataaacacagaaattacaaaaaaaatacatatgaattttttttttgtaatcttccttataaatatagaataatgaatcatataaaacatatcattattcatttatttacatttaaaattattgtttcagtatctttaatttattatgtatatataaaaataacttacaattttattaataaacaatatatgtttattaattcatgttttgtaatttatgggatagcgattttttttactgtctgtatttttcttttttaattatgttttaattgtattttatttttattattgttctttttatagtattattttaaaacaaaatgtattttctaagaacttataataataataaatataaattttaataaaaattatatttatcttttacaatatgaacataaagtacaacattaatatatagcttttaatatttttattcctaatcatgtaaatcttaaatttttctttttaaacatatgttaaatatttatttctcattatatataagaacatatttattaaatctagaattctatagtgagtcgtattacaattcactggccgtcgttttacaacgtcgtgactgggaaaaccctggcgttacccaacttaatcgccttgcagcacatccccctttcgccagctggcgtaatagcgaagaggcccgcaccgatcgcccttcccaacagttgcgcagcctgaatggcgaatggcgcctgatgcggtattttctccttacgcatctgtgcggtatttcacaccgcatatggtgcactctcagtacaatctgctctgatgccgcatagttaagccagccccgacacccgccaacacccgctgacgcgccctgacgggcttgtctgctcccggcatccgcttacagacaagctgtgaccgtctccgggagctgcatgtgtcagaggttttcaccgtcatcaccgaaacgcgcgagacgaaagggcctcgtgatacgcctatttttataggttaatgtcatgataataatggtttcttagacgtcaggtggcacttttcggggaaatgtgcgcggaacccctatttgtttatttttctaaatacattcaaatatgtatccgctcatgagacaataaccctgataaatgcttcaataatattgaaaaaggaagagtatgagtattcaacatttccgtgtcgcccttattcccttttttgcggcattttgccttcctgtttttgctcacccagaaacgctggtgaaagtaaaagatgctgaagatcagttgggtgcacgagtgggttacatcgaactggatctcaacagcggtaagatccttgagagttttcgccccgaagaacgttttccaatgatgagcacttttaaagttctgctatgtggcgcggtattatcccgtattgacgccgggcaagagcaactcggtcgccgcatacactattctcagaatgacttggttgagtactcaccagtcacagaaaagcatcttacggatggcatgacagtaagagaattatgcagtgctgccataaccatgagtgataacactgcggccaacttacttctgacaacgatcggaggaccgaaggagctaaccgcttttttgcacaacatgggggatcatgtaactcgccttgatcgttgggaaccggagctgaatgaagccataccaaacgacgagcgtgacaccacgatgcctgtagcaatgccaacaacgttgcgcaaactattaactggcgaactacttactctagcttcccggcaacaattaatagactggatggaggcggataaagttgcaggaccacttctgcgctcggcccttccggctggctggtttattgctgataaatctggagccggtgagcgtgggtctcgcggtatcattgcagcactggggccagatggtaagccctcccgtatcgtagttatctacacgacggggagtcaggcaactatggatgaacgaaatagacagatcgctgagataggtgcctcactgattaagcattggtaactgtcagaccaagtttactcatatatactttagattgatttaaaacttcatttttaatttaaaaggatctaggtgaagatcctttttgataatctcatgaccaaaatcccttaacgtgagttttcgttccactgagcgtcagaccccgtagaaaagatcaaaggatcttcttgagatcctttttttctgcgcgtaatctgctgcttgcaaacaaaaaaaccaccgctaccagcggtggtttgtttgccggatcaagagctaccaactctttttccgaaggtaactggcttcagcagagcgcagataccaaatactgtccttctagtgtagccgtagttaggccaccacttcaagaactctgtagcaccgcctacatacctcgctctgctaatcctgttaccagtggctgctgccagtggcgataagtcgtgtcttaccgggttggactcaagacgatagttaccggataaggcgcagcggtcgggctgaacggggggttcgtgcacacagcccagcttggagcgaacgacctacaccgaactgagatacctacagcgtgagctatgagaaagcgccacgcttcccgaagggagaaaggcggacaggtatccggtaagcggcagggtcggaacaggagagcgcacgagggagcttccagggggaaacgcctggtatctttatagtcctgtcgggtttcgccacctctgacttgagcgtcgatttttgtgatgctcgtcaggggggcggagcctatcgaaaaacgccagcaacgcggcctttttacggttcctggccttttgctggccttttgctcacatgttctttcctgcgttatcccctgattctgtggataaccgtattaccgcctttgagtgagctgataccgctcgccgcagccgaacgaccgagcgcagcgagtcagtgagcgaggaagcggaagagcgcccaatacgcaaaccgcctctccccgcgcgttggccgattcattaatgcagctggcacgacaggtttcccgactggaaagcgggcagtgagcgcaacgcaattaatgtgagttagctcactcattaggcaccccaggctttacactttatgcttccggctcgtatgttgtgtggaattgtgagcggataacaatttcacacaggaaacagctatgaccatgattacgccaag**ctatttaggtgacactatagaatactc**

Inserts:

>TryThrA-Ty1

GTTACTAGAGAAAAACTCGAATGGAAACATTGGGTAGAAATGAAAGAAAATATGAATATATATAATAAGTGGAAAAAATGGATAAAATGGAAGAAAAATAAATTAGCTAATTTTAATGAATGGTCAAAAAATTTTATAGAAAAATGGATACGAGAAAAACAATGGAACAATTGGATTAATGAAAGAAAAAAATATACATCTCAAAGAAAAAGTTTAGAACAACAATTTGGAGATAATATGGATAAAATGAACAAATTAAAAAAAAAAAAAATTTTGAAATTCTTCCCACTATTTAATTATAAAAGTGATTTGGAACCAATAATGGAAGAAGATGAAAACGAATACAACAGTTTTGATGAAAACGAAGAAGAAAACGATGAAAAAACAGGAGATGTTAATGTAGGAAAGACGGAAGCTTTGAATGTAGCAAAGACCGAAGGTTTGAATGTAGGAAAGACGGAAGATTTGAATGTAGCAAAGACAGAAGATTTGAATGTAGCAAAGACGGCAGATTTGAATGCAGAAAAAACAACAGATTTGAATTCAGAAAAAACAGCAGATTTGAATTCAGACAAAACAACAGATTTGAATCCAGAAAAAACAACAAATTTTAATACATACAAAGCAACAGATTTGAATGCAAACAAAACAGCAGATTTGAATTCAGACAAAACAACAGATTTGAATTCAGACAAAACAACAAATTTTAATACATACAGAACAACAGATTTGAATTCAGACAAAACAACAAATTTTAATACATACAAAACAGATTTGTATGCAGAAAAAACGACAGATGTTAATCTAGGAAAAACAACAAACCATAATGTAGCAAAAACAACAGATCAAAAGGTAGTAAAACATTCGTTAGATCACGAAGTAAGACAAATGATAGACCAAAAAGTAGCACAAATAATGAACCATGACTTAGAATCAACAGCAGAACAAAAAGCAGAAAAAAAAGGAGGAAAAGCAAAAGCGAAGACAAAAGTAAGAACAGTTGATGATGACGGAAATGAAATTAATGTTCCTAGGGAGGTGCACACCAACCAGGACCCCCTGGACGCCGAAGTCCATACAAATCAGGATCCTCTGGATGCCGAAGTGCACACCAATCAGGATCCCCTGGACGCT

>PTEF-Ty1

ATTATTATCCACATAATATGACATTTGGACAACAACAATATTTCCCATACTATAATCCATTAGAACAACAGAATTATCAACTACATCATATTATAAAACAACAGCAAAATTATCACCCACATCATATTATAAAACAACAGCAAAATCATAATCCACATCATATTTTACAAGAGCAAGAAAAACATCACCCACAAGGTATACCAAAGGAACAACCATATAATAATGTTCCTTATATCTTAAAAAAGGGGCTTGAACCCAAAACTCATAACCATGTAAAAGAAGATCAACCTAATATTAAACAGGGTGTTGTAAAGGGACAAGAACCACATGTTGATGATATGCATAACAATACAAAAGAACATAAGAATTTTAAAAATACAACCGATGTAAAACAACCAGCAAGTCATATATATAATAATTCATCAGAAAAACAAATTGAACATGTATATAATAAGTCTCCTGAAAAACAAATTGAACATGTATATAATAAGTCTCCTGAAAAACAAATTGAACATGTATATAATAATTCTCCTGAAAAACAAATTGAACATGTATATAATAATTCTCCTGAAAAACAAATTGAACATGTATATAATAATTCTCCTGAAAAACAAATTGAACATGTATATAATAATTCTCCTGAAAAACCTGCAAGACATACAAATAATATTTCATTAGAAAAACAAAATAGTCATAAATATAATGTTAATATACAAGATCGACATGATCCTGTATATTATAAATATGAAGATATGTTAAAAAGAGATAAAGATTTGTTTACAATTATCAACAACATTTGTGAACTGGAATTTAATTCTACAAATAACTATTTAATGAAAATAATTAATAACGACAAATTAAAACATAATTCTTTAAATGATAATGAAGCCATATTAAAAGAAATAACTAAAACTCAAAATGAATTGTTTTCTTTAAAATTACCATTAGAAATTAAAGTATCAATGGCTCTTCGTATAAGTGAACGTTTACGTGCCTTTGTTTTTGATAAAGATTTAACAGCTTATTATATAAAAAAATTAAAAGATATATTTAAATTAGAAACAGAGGCAGCAAAAAATTATTATTATTATGTAAAATGTCAAAAAACGTTTAGTGATAAAAAACGTTTGGTTAATAATTTAGATTCAATAAAATTATACTATGAATCACAAATTAATAAAAATTTTATTAGCATACCCAAAGATAAAATACCTACAGCTATATATCGTATATCCAACTTAGTTAATGATTTAATTTTTTTACTTCCACAATCAAATGCAAACAAAGCATTACCTAGGGAGGTGCACACCAACCAGGACCCCCTGGACGCCGAAGTCCATACAAATCAGGATCCTCTGGATGCCGAAGTGCACACCAATCAGGATCCCCTGGACGCT

>EMPIC3-Ty1

GCTTATCCTCTTTTAGAAGATGACTTAAGATCCATTAGGGTTGCTTTTGGAACTTGTCCTGGAAATAGCACAATGCAATTTGGAGAGAGATTTTGGCAAGGATTTTTTTTTGGTGTAATTATATTTTTTGTATTGTCTAAATATATACGTTCATATAAGAAGAGAAAAGGAATGgtaagaatataaaatatggaaatatgatattttacataaataaatataaatataaatataaataaatataaataaatataaataaatatatataaatataaataaatatatataaatataaatatatataaataaatataaataaatataaataaatatatataaatatatataaatataaataaatataaatatatatatataaatatatgtaaatataaataaatatgtataaatatataaatatatatatatattattaattttatagAGAATATTGAAATCTTCTTCCGTAACAAAAGATAGATTATATCGTGGAACCGTATTCGATGAAGATGATAATGTTGACGGCCACCATAAAAAGACCAAAGCCCATGGATTATTTGAAGAAATTGAAAAAAGAAATGGATTTGATTTAAATGATGATACAACAAACAGTAATAATTTGGGTGAAGGAACTCATAGTGCAGTGCATAATTTAGATGGACAAAAGGAAAGTCAATTATATGAAAACAATTCAGATGGAAGTACAAAAAATCACCTATCTGGAAGTAATTCAGATGGAAGGAATTCATCTCAACATAGCGTAGGTAGTCACTCTGTTAGTAGTCAAGAACATAAGAAAGAACACAGTACTCATGATTTACCTGCAGGAAGAAATTATTCTTTAGGAAATCTTAGCACAGGAACAACTTCACAAGGAAGTACATCATCAAGACATTACTCATTAGGAGGACAACCATCATCATCAGGACGTTCTTTTAGTGGTTCGAAATATAACACTAGCAATTTAGCTAGCTCAAGTACAACCGAAAGTAGCGTATCTGGTTTAAACACTAATGAAGCTCATGTA

>PeMP2-Ty1

AATAAAAAATCAATGCAAACTAAGAACTTTTTATCTGAAAGGAACTATGGAAGTATAGATCAAAATGTGAGGACTAAAAATAAAAGAAGATTAATGAAATTCCAAAGTAAGAGTAAAGCAAAATCGTTCCTTTTTTTATTGGAACTTATGGTATTCTCCCTTTTCATATGGATTTTAAAGAGTGCAAAGCATgtaagtttatattttctgtatatatatatatatatatatatatatatatatatatatatatatttatttatttatttatttattgatttattcatgtacacctatatttgtattgctctacataaaacgataaaaaattatgttatccctacataaaaataatgattttgtatttctatccataataactacctatatataaacacatatttctttatttatccctttacattaatttcttagAACGTATCATCCAAATCGATATATAATAAAAATAAATTCCATAACACGTTCAATAGAAGAGATACAAGAGTTTTAGCAGAGCAAGAAGATCAATACATAAGGAACCCAAATAATTCTAATTATCCTGATAGAGACCTTGACATCTGTAATTGGGATGAACCTCATAATCCTGAAAAAAATCCTTGTGCCATTCCACAAGACGATTTATCCAATGAAGGCATAGAAATAATTGATTATGTTGATAAAAAAGAAGAAGAGAAACTTAAAAATGATTTGAAAATACATTTTGGCAAACAACACTATGCATTAATTAAGGATATATATAAACCTATACAATCTTATGCAAACAATTTTTCAAAGGTCGTAGATCATTATGTTGGTAGTTTAAAAAGAATTAACAATGCATTCAATTTAGATATTTTACGTACTGTTATATGCACAGCATTCCTTATACAACCAATAGTTTTATTAATGAAACTTCACCCTGAACTTCTTGTTCACATACCAATCAATATTATAATATCCATGGTTGTATTTAAATTAATGCATAAATCACTACGTAACGAACAAGAATCAAATAAACCTAGGGAGGTGCACACCAACCAGGACCCCCTGGACGCCGAAGTCCATACAAATCAGGATCCTCTGGATGCCGAAGTGCACACCAATCAGGATCCCCTGGACGCT

Inserts:

>PTP1-TGD

GTGAATAAAGATAATAGGAAAATTCATAAGGCACATATTAGTAGATATGTAGGATATAGGAAACGAGAAATGAATAATACTTTTTTTAGCAAATCCTATAAGTTGTTTTTTTCAAATAATAAATATGAATATTCAAAAGATGAAGAAAAAAAAAATATATATCCTTTATCATTAAAACTTTTAATATTAACAGTTTTTATATGTATATTGCAATTTTCTAATAATAATgtaagacatttgtacataaaaaaataaatagaggacttttatcatcatattaattattataaatatgtacatttttttacatatatttatggaattatatgtttaaacttttctttgtgtttatatatttttatagtcttatatatttatatgtttatataattatatgattacatatttataagtttagacaatttatatgtttattattttttatttttttttttttattttttatatatttttttattttaaatttttagGGATATTTCGAAAAGTATGTTAATAAAAATTCAGGAAATGACAATGAGTATTATTTTAATATAAAATATAATAGATCCTTAGCAGAATTAAAAAAAAGAATTGGCGCTATAAGAGAAATGATAAATAACGATTCCTTTGGTACC

>TryThrA-TGD

AATTTAGAGCAGTTTAAAAATATAAACAAAGATCTGGCTACAAATCTTTTTTCTCAACTATCATTTTTAAAAAATGAGAATAAGTTTTTGAGCCAGGGTAAATCTCTTATTAAATTTCTTATCGGAATAGCCATATTTCTAGTCGTTCTCATATTTATAAAATCGTCTCATCCAgtaagtttattatatatatatatatatatatatataaattctttttagtacatttttaatattttctctttgtatttaaaaaaaaaaaaaaatacatatacattaatatatatatatatatatatatatatatatatatatttttttctctctctctcttcctttttagGCTCTCAAAGAGAAGAAGAAAAAGGTGCTAGAATTTTTTGAAAACCTTGTATTGAATAAGAAAAAAAAAGAGAATATAACAGCAGCGATCGCATCCAAAGAATTGGCTGATGCTGAAACAACAGATACAAGCGATTCAGAAGATGAAGATCATATTATTAATAAAAAAGTGAAAAGAAGAAAAAGGAATATCATAAATAATCCTGATGAGAAAGTACATAACGTAAAAGAAAAAAATAGTAAAAGTAAAAATGAAGAAGATAAAACAGATGAAAGTTACAATGAAACTAGCTTATTATCTTCTGATGAAGAAGGTGAGGTGAATTTAGAGGATTGGAAAAAAAACGAATGGATTAAATGGATGGATGAAACAGAAGAAGAATGGCAACTTCTAAAATTATGGTTGGAAGGAGAAAAAAATAAATGGCTTGAAGGGAAAAATAAGGAATATGATATATGGTTAAACCATATGAATAGTAAATGGACAAATTATAATAAAGATATAGATGAAGAATATGATTCAAATGTTTTTAAAGATTCTTACAAATGGAATGAAAAACAATGGGAACAATGGATGAAAACCGAAGGAAAAGAATTCATGTTACAAGATTTTAAAAGATGGTTAGAAGACAGTGAAGGATATTTAAAATCATGGCGTACGATGAGTAAAGGAGAAGAACTTTTCACTGGAGTTGTCCCAATTCTTGTTGAATTAGATGGTGATGTTAATGGGCACAAATTTTCTGTCAGTGGAGAGGGTGAAGGTGATGCAACATACGGAAAACTTACCCTTAAATTTATTTGCACTACTGGAAAACTACCTGTTCCATGGCCAACACTTGTCACTACTTTCGCGTATGGTCTTCAATGCTTTGCGAGATACCCAGATCATATGAAACAGCATGACTTTTTCAAGAGTGCCATGCCCGAAGGTTATGTACAGGAAAGAACTATATTTTTCAAAGATGACGGGAACTACAAGACACGTGCTGAAGTCAAGTTTGAAGGTGATACCCTTGTTAATAGAATCGAGTTAAAAGGTATTGATTTTAAAGAAGATGGAAACATTCTTGGACACAAATTGGAATACAACTATAACTCACACAATGTATACATCATGGCAGACAAACAAAAGAATGGAATCAAAGTTAACTTCAAAATTAGACACAACATTGAAGATGGAAGCGTTCAACTAGCAGACCATTATCAACAAAATACTCCAATTGGCGATGGCCCTGTCCTTTTACCAGACAACCATTACCTGTCCACACAATCTGCCCTTTCGAAAGATCCCAACGAAAAGAGAGACCACATGGTCCTTCTTGAGTTTGTAACAGCTGCTGGGATTACACATGGCATGGATGAGCTCTACAAA

>PTEF-TGD

TTAAAGAAATATATTATATTAATATATATCGGTGTAATTCTAAATTTCATAACTAAAAATAATAATGTAGTGTCTGTTCCTGAGCCCTTTTTATCACAAAACAAAGATTCTTTTGAAGAAAAAAAATATACGTATGGGGATAATTTACAATTGGGGGCATCAACTATAAATACCCCTAAAACACAATCACAAGAAAATAAAGATATAAATAAAGAAACAAAAAATACAATAATAAAAAAAACGAATAATTTTCCAAGTACCTTAAATGAGAAATTTCCTCATAAAATTCAATTAACCAATAAAGAAAATAAAGAAGATGAACAAAATAAAGAAAATAAAAAAGATGAACAAAATAAAGAAGATGAACAAAATAAACAAAATAAAGAAGATGAACAAAATAAACAAAACAAGGATAAAAAAAATATAGTAAGTAACAAATTATCGGGAAACAATGAACAACAGAATAATTCTATTCCAAAATCAATACAAAAACCAGAGAATTGTGTCAAAAAACAAAGCAATCAATTTCCTCGTAGTTATCCAGAATTCTTTGAAGCAAATTTTGGACCCATAGATGAACTTATGGATGAAACGGATTATAGTAGTGATGATTTAGAAGATCAACTAAATTATCGTACGATGAGTAAAGGAGAAGAACTTTTCACTGGAGTTGTCCCAATTCTTGTTGAATTAGATGGTGATGTTAATGGGCACAAATTTTCTGTCAGTGGAGAGGGTGAAGGTGATGCAACATACGGAAAACTTACCCTTAAATTTATTTGCACTACTGGAAAACTACCTGTTCCATGGCCAACACTTGTCACTACTTTCGCGTATGGTCTTCAATGCTTTGCGAGATACCCAGATCATATGAAACAGCATGACTTTTTCAAGAGTGCCATGCCCGAAGGTTATGTACAGGAAAGAACTATATTTTTCAAAGATGACGGGAACTACAAGACACGTGCTGAAGTCAAGTTTGAAGGTGATACCCTTGTTAATAGAATCGAGTTAAAAGGTATTGATTTTAAAGAAGATGGAAACATTCTTGGACACAAATTGGAATACAACTATAACTCACACAATGTATACATCATGGCAGACAAACAAAAGAATGGAATCAAAGTTAACTTCAAAATTAGACACAACATTGAAGATGGAAGCGTTCAACTAGCAGACCATTATCAACAAAATACTCCAATTGGCGATGGCCCTGTCCTTTTACCAGACAACCATTACCTGTCCACACAATCTGCCCTTTCGAAAGATCCCAACGAAAAGAGAGACCACATGGTCCTTCTTGAGTTTGTAACAGCTGCTGGGATTACACATGGCATGGATGAGCTCTACAAA

>EMPIC3-TGD

GCTTATCCTCTTTTAGAAGATGACTTAAGATCCATTAGGGTTGCTTTTGGAACTTGTCCTGGAAATAGCACAATGCAATTTGGAGAGAGATTTTGGCAAGGATTTTTTTTTGGTGTAATTATATTTTTTGTATTGTCTAAATATATACGTTCATATAAGAAGAGAAAAGGAATGgtaagaatataaaatatggaaatatgatattttacataaataaatataaatataaatataaataaatataaataaatataaataaatatatataaatataaataaatatatataaatataaatatatataaataaatataaataaatataaataaatatatataaatatatataaatataaataaatataaatatatatatataaatatatgtaaatataaataaatatgtataaatatataaatatatatatatattattaattttatagAGAATATTGAAATCTTCTTCCGTAACAAAAGATAGATTATATCGTGGAACCGTATTCGATGAAGATGATAATGTTGACGGCCACCATAAAAAGACCAAAGCCCATGGATTATTTGAAGAAATTGAAAAAAGAAATCGTACGATGAGTAAAGGAGAAGAACTTTTCACTGGAGTTGTCCCAATTCTTGTTGAATTAGATGGTGATGTTAATGGGCACAAATTTTCTGTCAGTGGAGAGGGTGAAGGTGATGCAACATACGGAAAACTTACCCTTAAATTTATTTGCACTACTGGAAAACTACCTGTTCCATGGCCAACACTTGTCACTACTTTCGCGTATGGTCTTCAATGCTTTGCGAGATACCCAGATCATATGAAACAGCATGACTTTTTCAAGAGTGCCATGCCCGAAGGTTATGTACAGGAAAGAACTATATTTTTCAAAGATGACGGGAACTACAAGACACGTGCTGAAGTCAAGTTTGAAGGTGATACCCTTGTTAATAGAATCGAGTTAAAAGGTATTGATTTTAAAGAAGATGGAAACATTCTTGGACACAAATTGGAATACAACTATAACTCACACAATGTATACATCATGGCAGACAAACAAAAGAATGGAATCAAAGTTAACTTCAAAATTAGACACAACATTGAAGATGGAAGCGTTCAACTAGCAGACCATTATCAACAAAATACTCCAATTGGCGATGGCCCTGTCCTTTTACCAGACAACCATTACCTGTCCACACAATCTGCCCTTTCGAAAGATCCCAACGAAAAGAGAGACCACATGGTCCTTCTTGAGTTTGTAACAGCTGCTGGGATTACACATGGCATGGATGAGCTCTACAAA

>PeMP2-TGD

AATAAAAAATCAATGCAAACTAAGAACTTTTTATCTGAAAGGAACTATGGAAGTATAGATCAAAATGTGAGGACTAAAAATAAAAGAAGATTAATGAAATTCCAAAGTAAGAGTAAAGCAAAATCGTTCCTTTTTTTATTGGAACTTATGGTATTCTCCCTTTTCATATGGATTTTAAAGAGTGCAAAGCATgtaagtttatattttctgtatatatatatatatatatatatatatatatatatatatatatatttatttatttatttatttattgatttattcatgtacacctatatttgtattgctctacataaaacgataaaaaattatgttatccctacataaaaataatgattttgtatttctatccataataactacctatatataaacacatatttctttatttatccctttacattaatttcttagAACGTATCATCCAAATCGATATATAATAAAAATAAATTCCATAACACGTTCAATAGAAGAGATACAAGAGTTTTAGCAGAGCAAGAAGATCAATACATAAGGAACCCAAATAATTCTAATTATCCTGATAGAGACCTTGACATCTGTAATCGTACGATGAGTAAAGGAGAAGAACTTTTCACTGGAGTTGTCCCAATTCTTGTTGAATTAGATGGTGATGTTAATGGGCACAAATTTTCTGTCAGTGGAGAGGGTGAAGGTGATGCAACATACGGAAAACTTACCCTTAAATTTATTTGCACTACTGGAAAACTACCTGTTCCATGGCCAACACTTGTCACTACTTTCGCGTATGGTCTTCAATGCTTTGCGAGATACCCAGATCATATGAAACAGCATGACTTTTTCAAGAGTGCCATGCCCGAAGGTTATGTACAGGAAAGAACTATATTTTTCAAAGATGACGGGAACTACAAGACACGTGCTGAAGTCAAGTTTGAAGGTGATACCCTTGTTAATAGAATCGAGTTAAAAGGTATTGATTTTAAAGAAGATGGAAACATTCTTGGACACAAATTGGAATACAACTATAACTCACACAATGTATACATCATGGCAGACAAACAAAAGAATGGAATCAAAGTTAACTTCAAAATTAGACACAACATTGAAGATGGAAGCGTTCAACTAGCAGACCATTATCAACAAAATACTCCAATTGGCGATGGCCCTGTCCTTTTACCAGACAACCATTACCTGTCCACACAATCTGCCCTTTCGAAAGATCCCAACGAAAAGAGAGACCACATGGTCCTTCTTGAGTTTGTAACAGCTGCTGGGATTACACATGGCATGGATGAGCTCTACAAA

>PTP7-TGD

GCAAAAGATAGTCAAAAGAACTTGAATGTTTCTAATAATAATAACGTCCAATGCACCATGGGAAGATCAAGCCAGAACATAAATAAATCCGATTCAAAAGGAAAAATAAAAAGGTGCACTTATGCCTATAAAATTTTATTATGTACAATTTTTATCTGGATATGTCAATGTTTTTATAATgtaagtttatatatatatatatatatatatatatataaatatatttttttttttgtaagatatttatatgtaatatatacattttattaatttgcatatatgtatttatttatttgttattttttttgtgaatatatgctttgtgtattcctatattctttttatatcaattttttttatttcacatatttattgtatttctttttttttttatgaatattacattctagttttttattcccaaaattaaatatgtgcatgtaaatattcttaacaataatatgatatatatatatatgtatatatatatgccattgaagtgatattcttatgtacattttaattttaatatatattttaattttttttttttttttttttttgatatatagAAATCATATTATGTATATAAAAAAGACGGAAGAAGAAATAAAGGAAAGAAAATATTAGGTATAAGAATTAATAAATCCTTAGCCGAAATGGATCATACAAAATATCACCCAGAATATTATGATGAAGTTCAAGAAAATTATGACCCTTATCGTACGATGAGTAAAGGAGAAGAACTTTTCACTGGAGTTGTCCCAATTCTTGTTGAATTAGATGGTGATGTTAATGGGCACAAATTTTCTGTCAGTGGAGAGGGTGAAGGTGATGCAACATACGGAAAACTTACCCTTAAATTTATTTGCACTACTGGAAAACTACCTGTTCCATGGCCAACACTTGTCACTACTTTCGCGTATGGTCTTCAATGCTTTGCGAGATACCCAGATCATATGAAACAGCATGACTTTTTCAAGAGTGCCATGCCCGAAGGTTATGTACAGGAAAGAACTATATTTTTCAAAGATGACGGGAACTACAAGACACGTGCTGAAGTCAAGTTTGAAGGTGATACCCTTGTTAATAGAATCGAGTTAAAAGGTATTGATTTTAAAGAAGATGGAAACATTCTTGGACACAAATTGGAATACAACTATAACTCACACAATGTATACATCATGGCAGACAAACAAAAGAATGGAATCAAAGTTAACTTCAAAATTAGACACAACATTGAAGATGGAAGCGTTCAACTAGCAGACCATTATCAACAAAATACTCCAATTGGCGATGGCCCTGTCCTTTTACCAGACAACCATTACCTGTCCACACAATCTGCCCTTTCGAAAGATCCCAACGAAAAGAGAGACCACATGGTCCTTCTTGAGTTTGTAACAGCTGCTGGGATTACACATGGCATGGATGAGCTCTACAAA

**>pFNT_SBP1-mCherry**

xxxxx= Nmd3 Promotor

xxxxx= SBP1

xxxxx= mCherry

xxxxx= Cam Promotor

xxxxx = FNT (mutation conferring resistance is underlined, leading to G 107 S aa change)

CGGCCGCTAACGTAACAGACTTAGGAGGAGATCTttattattattacatGTTGAAATATAAATTTCAAAAAAAATGATCACAAAATATACACTTAAATATAGGTACAATAAAAAAAAAAAATAAAAATATAATTACAAGATAATATTTTTTCCTGCTATCAAATTTTTATATATTCTCCTCAAAGAAAAAATATAAATAAATGAAGTAAATTAAAAAAAAAATTTTCTTTTTCTTCTTCTTTTGTAATTCCTTATTTATACATATTTTACTATATTTTCATAAAAATAAATTGTCATATTATATAAATATATATACCAAACCATAATTATATAGCCCTCAACATATTTTTATGATGTTTTTTCTTTTTAAATGTGATACGTAATTAAATATAATAATATATATTAAATATTATATTTTGTAATATTTACTTTCATGAGGTTTAATAAATTATAAAGAGAGAATAAAAAAAAAAAAAAAAAAAATTTATCCATATACAAATTATTAATTTATTTTTATTTTTTATTTCCCTTTGTATATATTATAAAAAAATAATCTACATAATTTTATATGATGATAATTACAATATAATATTTATAATATATATTATTTGTTAAGAGAAAAAAAAAATAAATATATACCTTCTTTTTGAGAATTGAATAAATTGTTTAATATATATATATATATATATAAATATATTATACATTTATGGTGAAAAAAAATGTTGTATTTAATTAGATTTAATATATATATATAAAAAATAGCGTATTTAAAATAATATATATATATATATATTATTATTATTACAAAATGACGGATATTATAAAAGTATATATCTATATATATGTATATATATATAATATTATTTTACTATATATATATAATATATAAATGTAAATGCATATAGTATCTATGTATATTATATATATATATAATATTAATTACATATCTAAGTTTTCTTTTCTTTTCTTTTTTTTTTTTTTTTTTTTTATTTTTTATAGAGAGCCCGTTATATATTATATATTAAAATTTTTTATAATAGCATTTATATACAATATTTTATTAACTAAAAGAAAAAAAAAAAAAAAAAAAAAAAAAAAACGAGGAAATTTATATTTCTTTAACAACATTTTAATTAAAATCATGATAATACAATTTCAAATCATTTTGTAATTATATAAATAAATATATATATATATATATATATATATATGTACTTTTAAATTAGGAATATTCTCATTTATAAATATATCTTATTTTTTAAATTGGTATAAAAAAAAAAAAAAAAATAAGAAACCGTTGATTAAATAATACATATATAATATAAATATATTTTATAAATATATATTTATATATATATATATATATATTTATAACGTATATCATTTTAAAGATAActcgagATGTGTAGCGCAGCTCGAGCATTTGATTTTTTTACTGATTTAGCCGACGAACCAACACAATTACAGGATGCAGTACCAGAGACAACCGAAAAATTGGCCGAAGTAGTTTCGGATGCAGCAACAAATGTTACTGATGCAGTAAGTGATACAGCTAGTGGTATTGGAAGTTTAGTTGGAGAAGCAGCTAGTAGTTTAGGAAATTTAGTTGGTGAAGCAGCAAGCGGTATAGGAAATATAGTTGGAGGTGCAGCAAGCGGTATAGGAAATATAGTTGGAGGTGCAGCAAGCGGTATAGGAAGTTTAGTTGGTGATGCAGCAAGCGGTTTAGGAAATTTAGTTGGTGATGCAGCAGAGGCACTTGCAACTACCGAATTAAAAGATGTAATACCAGAAAATACTGAATCCACAACTGATTTGGTACCATCTGAGGTATCACCTCCAGTAGATGATTATCTCGACGATGACGGTTTTTCAAGCTTTAGAGAATTTCTTGAAAGTACTCCTTGTTGGCAACGTAGAATGGCTCAAGAAGCTTTACTTAATGAATACGAAGTAGAATCTCCAGCCGAATCTATGTCCCCTATTCTTAGAGTACAATTTTTTGCAGATTTTGCAAAACAAGCCGTACATGTTGCTAAACAAAATTATCTCTATGTTGTGATATTCCTATTCTTTGTTATTAACATATTATTGTTCATCAACTTTTACAACTTAGGAAAAAGAAAAGGATATTACCTAGCAAAAAAACAAAAAAAAGAACAAATGCTAGAACAAAACCCAGAACAAAACCCAGAACAAAACGCACAACAAAACGCACAACAAAACGCACAACAAAACGCACAACAAAACGCACAACAAAACGCACAACAAAACGCACAACAAAACACACAACAAAACACACAACAAAAAACACAACAGAACCCACAACAAAACGCACAACAAAACACACAACAAAACACACAACAACAATCCACAACCAAATCCACAACAAAAACAGTTGCTAGAGAAACCggtaccATGGTGAGCAAGGGCGAGGAGGATAACATGGCCATCATCAAGGAGTTCATGCGCTTCAAGGTGCACATGGAGGGCTCCGTGAACGGCCACGAGTTCGAGATCGAGGGCGAGGGCGAGGGCCGCCCCTACGAGGGCACCCAGACCGCCAAGCTGAAGGTGACCAAGGGTGGCCCCCTGCCCTTCGCCTGGGACATCCTGTCCCCTCAGTTCATGTACGGCTCCAAGGCCTACGTGAAGCACCCCGCCGACATCCCCGACTACTTGAAGCTGTCCTTCCCCGAGGGCTTCAAGTGGGAGCGCGTGATGAACTTCGAGGACGGCGGCGTGGTGACCGTGACCCAGGACTCCTCCCTGCAGGACGGCGAGTTCATCTACAAGGTGAAGCTGCGCGGCACCAACTTCCCCTCCGACGGCCCCGTAATGCAGAAGAAGACCATGGGCTGGGAGGCCTCCTCCGAGCGGATGTACCCCGAGGACGGCGCCCTGAAGGGCGAGATCAAGCAGAGGCTGAAGCTGAAGGACGGCGGCCACTACGACGCTGAGGTCAAGACCACCTACAAGGCCAAGAAGCCCGTGCAGCTGCCCGGCGCCTACAACGTCAACATCAAGTTGGACATCACCTCCCACAACGAGGACTACACCATCGTGGAACAGTACGAACGCGCCGAGGGCCGCCACTCCACCGGCGGCATGGACGAGCTGTACAAGTAACCCGGGTCGAGGGATATGGCAGCTTAATGTTCGTTTTTCTTATTTATATATTTATACCAATTGATTGTATTTATAACTGTAAAAATGTGTATGTTGTGTGCATATTTTTTTTTGTGCATGCACATGCATGTAAATAGCTAAAATTATGAACATTTTATTTTTTGTTCAGAAAAAAAAAACTTTACACACATAAAATGGCTAGTATGAATAGCCATATTTTATATAAATTAAATCCTATGAATTTATGACCATATTAAAAATTTAGATATTTATGGAACATAATATGTTTGAAACAATAAGACAAAATTATTATTATTATTATTATTTTTACTGTTATAATTATGTGTCTCCTTCAATGATTCATAAATAGTTGGACTTGATTTTTAAAATGTTTATAATATGATTAGCATAGTTAAATAAAAAAAGTTGAAAAATTAAAAAAAAACATATAAACACAAATGATGGTTTTTCCTTCAATTTCGATATCAATTTATAGAAACAAAATATATACTTGTATAATTTTATTTTTTTATATAAATCATTACATATATAATTATACAATATTTTTTCTAAGAGATAATTATATATTAATATATATAAAAAAAGGTGTTTTTTTTTTTTTTTTTTATTTTTATTTTTATTTTATGGTAATATTTTATTTTCCTTATTTTATAAATTATATTAGTTTATATGTGATTAATTTTATATATTATCAATTTATATATTTTTAAATGCTTACTTAATTATCTTTTTTTTTTTTTTTTTTTTTTTTTCCCCTCTTTTTATATTAATTTATTTTTGAAAAAATTGATATATATATATATATATAATATATATATATACATGTAGTAGTATTAAACAATGTATAATATATATAAATAATATATTTATATATTTCATTTCAATTTTAATTTTTTTTGGTTTTTTTTTTTTTTCTTTTTGTCATATTTAAAAAAAATTATATTCATATAAGTTATGCATTTTTTATAAACATTATTCAATATATGTATAATATAATATATATATATATATTAATGTATTATTCCAATGTGCATGATAAAAGAAAAAAATAATATTTATAAAAAAAAAGAAAAATAAAACAAAAAAAGAAAAAAAAAAAAAAAAAAAAAAAAATACAAAAATAAATAATATAATTTATAATTATATATTCTTGTCACAATAAAAATATATATATATATATATATATTTATAATATGTATATTTTAAACTAGAAAAGGAATAACTAATATTTTATTTATTATCATTCAAGATTTATATTTTATAATAATAAATACCTAATAGAAATATATCAGGATCCATGCCTCCCAACAACTCGAAATATGTTCTTGACCCTGTTTCTATCAAAAGCGTGTGCGGAGGGGAAGAAAGCTATATCCGTTGTGTTGAGTATGGCAAGAAGAAAGCCCATTACAGTAATCTGAATCTTTTAGCCAAAGCTATTCTGGCAGGGATGTTCGTAGGCCTGTGTGCACATGCGTCAGGAATCGCGGGTGGGTTGTTCTACTACCACAAACTGCGTGAGATTGTTGGAGCGTCTATGAGTGTCTTTGTGTACGGCTTTACATTTCCGATAGCATTCATGTGCATCATCTGTACCGGTTCGGATTTGTTTACC**A**GTAATACACTAGCGGTCACAATGGCACTGTACGAAAAGAAAGTTAAACTTTTAGACTATCTCCGGGTTATGACCATCTCGTTATTTGGGAACTATGTCGGTGCAGTCTCATTTGCTTTCTTTGTGTCCTACTTGTCGGGTGCTTTCACTAACGTGCATGCTGTGGAAAAGAATCATTTCTTTCAGTTTCTGAATGATATCGCAGAGAAGAAAGTTCACCATACATTTGTGGAATGCGTTAGCTTAGCCGTGGGTTGTAACATATTTGTATGCTTGGCGGTGTATTTCGTGCTGACTCTCAAAGATGGGGCTGGGTATGTATTTTCTGTGTTCTTTGCAGTTTATGCCTTTGCGATTGCCGGCTATGAGCATATCATAGCCAACATCTATACCCTGAATATAGCCCTGATGGTGAATACGAAGATTACGGTGTATCAGGCCTATATTAAAAACCTGCTGCCGACACTGTTGGGCAACTACATTGCGGGTGCGATTGTCCTGGGACTGCCGCTGTATTTCATTTACAAAGAACACTACTACAACTTCGAACGCAGTAAACGCGACAACAATGATGCGCAAATGAAATCCCTGAGCATTGAACTGCGCAACAAGCTTATTTAATAATAGATTAAAAATATTATAAAAATAAAAACATAAACACAGAAATTACAAAAAAAATACATATGAATTTTTTTTTGTAATCTTCCTTATAAATATAGAATAATGAATCATATAAAACATATCATTATTCATTTATTTACATTTAAAATTATTGTTTCAGTATCTTTAATTTATTATGTATATATAAAAATAACTTACAATTTTATTAATAAGCAATATATGTTTATTAATTCATGTTTTGTAATTTATGGGATAGCGATTTTTTTTACTGTCTGTATTTTTCTTTTTTAATTATGTTTTAATTGTATTTTATTTTTATTATTGTTCTTTTTATAGTATTATTTTAAAACAAAATGTATTTTCTAAGAACTTATAATAATAATAAATATAAATTTTAATAAAAATTATATTTATCTTTTACAATATGAACATAAAGTACAACATTAATATATAGCTTTTAATATTTTTATTCCTAATCATGTAAATCTTAAATTTTTCTTTTTAAACATATGTTAAATATTTATTTCTCATTATATATAAGAACATATTTATTAAATCTAGAATTCTATAGTGAGTCGTATTACAATTCACTGGCCGTCGTTTTACAACGTCGTGACTGGGAAAACCCTGGCGTTACCCAACTTAATCGCCTTGCAGCACATCCCCCTTTCGCCAGCTGGCGTAATAGCGAAGAGGCCCGCACCGATCGCCCTTCCCAACAGTTGCGCAGCCTGAATGGCGAATGGCGCCTGATGCGGTATTTTCTCCTTACGCATCTGTGCGGTATTTCACACCGCATATGGTGCACTCTCAGTACAATCTGCTCTGATGCCGCATAGTTAAGCCAGCCCCGACACCCGCCAACACCCGCTGACGCGCCCTGACGGGCTTGTCTGCTCCCGGCATCCGCTTACAGACAAGCTGTGACCGTCTCCGGGAGCTGCATGTGTCAGAGGTTTTCACCGTCATCACCGAAACGCGCGAGACGAAAGGGCCTCGTGATACGCCTATTTTTATAGGTTAATGTCATGATAATAATGGTTTCTTAGACGTCAGGTGGCACTTTTCGGGGAAATGTGCGCGGAACCCCTATTTGTTTATTTTTCTAAATACATTCAAATATGTATCCGCTCATGAGACAATAACCCTGATAAATGCTTCAATAATATTGAAAAAGGAAGAGTATGAGTATTCAACATTTCCGTGTCGCCCTTATTCCCTTTTTTGCGGCATTTTGCCTTCCTGTTTTTGCTCACCCAGAAACGCTGGTGAAAGTAAAAGATGCTGAAGATCAGTTGGGTGCACGAGTGGGTTACATCGAACTGGATCTCAACAGCGGTAAGATCCTTGAGAGTTTTCGCCCCGAAGAACGTTTTCCAATGATGAGCACTTTTAAAGTTCTGCTATGTGGCGCGGTATTATCCCGTATTGACGCCGGGCAAGAGCAACTCGGTCGCCGCATACACTATTCTCAGAATGACTTGGTTGAGTACTCACCAGTCACAGAAAAGCATCTTACGGATGGCATGACAGTAAGAGAATTATGCAGTGCTGCCATAACCATGAGTGATAACACTGCGGCCAACTTACTTCTGACAACGATCGGAGGACCGAAGGAGCTAACCGCTTTTTTGCACAACATGGGGGATCATGTAACTCGCCTTGATCGTTGGGAACCGGAGCTGAATGAAGCCATACCAAACGACGAGCGTGACACCACGATGCCTGTAGCAATGCCAACAACGTTGCGCAAACTATTAACTGGCGAACTACTTACTCTAGCTTCCCGGCAACAATTAATAGACTGGATGGAGGCGGATAAAGTTGCAGGACCACTTCTGCGCTCGGCCCTTCCGGCTGGCTGGTTTATTGCTGATAAATCTGGAGCCGGTGAGCGTGGGTCTCGCGGTATCATTGCAGCACTGGGGCCAGATGGTAAGCCCTCCCGTATCGTAGTTATCTACACGACGGGGAGTCAGGCAACTATGGATGAACGAAATAGACAGATCGCTGAGATAGGTGCCTCACTGATTAAGCATTGGTAACTGTCAGACCAAGTTTACTCATATATACTTTAGATTGATTTAAAACTTCATTTTTAATTTAAAAGGATCTAGGTGAAGATCCTTTTTGATAATCTCATGACCAAAATCCCTTAACGTGAGTTTTCGTTCCACTGAGCGTCAGACCCCGTAGAAAAGATCAAAGGATCTTCTTGAGATCCTTTTTTTCTGCGCGTAATCTGCTGCTTGCAAACAAAAAAACCACCGCTACCAGCGGTGGTTTGTTTGCCGGATCAAGAGCTACCAACTCTTTTTCCGAAGGTAACTGGCTTCAGCAGAGCGCAGATACCAAATACTGTCCTTCTAGTGTAGCCGTAGTTAGGCCACCACTTCAAGAACTCTGTAGCACCGCCTACATACCTCGCTCTGCTAATCCTGTTACCAGTGGCTGCTGCCAGTGGCGATAAGTCGTGTCTTACCGGGTTGGACTCAAGACGATAGTTACCGGATAAGGCGCAGCGGTCGGGCTGAACGGGGGGTTCGTGCACACAGCCCAGCTTGGAGCGAACGACCTACACCGAACTGAGATACCTACAGCGTGAGCTATGAGAAAGCGCCACGCTTCCCGAAGGGAGAAAGGCGGACAGGTATCCGGTAAGCGGCAGGGTCGGAACAGGAGAGCGCACGAGGGAGCTTCCAGGGGGAAACGCCTGGTATCTTTATAGTCCTGTCGGGTTTCGCCACCTCTGACTTGAGCGTCGATTTTTGTGATGCTCGTCAGGGGGGCGGAGCCTATCGAAAAACGCCAGCAACGCGGCCTTTTTACGGTTCCTGGCCTTTTGCTGGCCTTTTGCTCACATGTTCTTTCCTGCGTTATCCCCTGATTCTGTGGATAACCGTATTACCGCCTTTGAGTGAGCTGATACCGCTCGCCGCAGCCGAACGACCGAGCGCAGCGAGTCAGTGAGCGAGGAAGCGGAAGAGCGCCCAATACGCAAACCGCCTCTCCCCGCGCGTTGGCCGATTCATTAATGCAGCTGGCACGACAGGTTTCCCGACTGGAAAGCGGGCAGTGAGCGCAACGCAATTAATGTGAGTTAGCTCACTCATTAGGCACCCCAGGCTTTACACTTTATGCTTCCGGCTCGTATGTTGTGTGGAATTGTGAGCGGATAACAATTTCACACAGGAAACAGCTATGACCATGATTACGCCAAGCTATTTAGGTGACACTATAGAATACTC
